# SARS-CoV-2 spike RBD and nucleocapsid encoding DNA vaccine elicits T cell and neutralising antibody responses that cross react with variants

**DOI:** 10.1101/2021.06.18.448932

**Authors:** VA Brentville, M Vankemmelbeke, RL Metheringham, P Symonds, KW Cook, RA Urbanowicz, T Tsoleridis, CM Coleman, K-C Chang, A Skinner, E Dubinina, I Daniels, S Shah, M Argonza, J Delgado, V Dwivedi, V Kulkarni, JE Dixon, AG Pockley, SE Adams, SJ Paston, JM Daly, JK Ball, LG Durrant

**Affiliations:** Scancell Ltd, The University of Nottingham Biodiscovery Institute, Nottingham, UK; Wolfson Centre for Global Virus Research, The University of Nottingham, Queen’s Medical Centre, Derby Road, Nottingham, UK; NIHR Nottingham Biomedical Research Centre, Nottingham University Hospitals NHS Trust and the University of Nottingham; Queen’s Medical Centre, Derby Road, Nottingham, UK; School of Life Sciences, The University of Nottingham, Queen’s Medical Centre, Derby Road, Nottingham, UK; Department of Infection Biology and Microbiomes, Institute of Infection, Veterinary and Ecological Sciences, University of Liverpool, Liverpool, UK; School of Veterinary Medicine and Science, University of Nottingham, Nottingham, UK; Division of Regenerative Medicine & Cellular Therapies (RMCT), The University of Nottingham Biodiscovery Institute, School of Pharmacy, University Park, Nottingham, UK; John van Geest Cancer Research Centre & Centre for Health, Ageing and Understanding Disease (CHAUD), School of Science and Technology, Nottingham Trent University, Clifton campus, Nottingham, UK; Scancell Ltd, Bellhouse Building, Sanders Road, Oxford Science Park, Oxford, OX4 4GD, UK; Division of Cancer and Stem Cells, The University of Nottingham Biodiscovery Institute, Nottingham UK; Disease Intervention & Prevention Program, Texas Biomedical Research Institute, San Antonio, Texas 78227, USA

## Abstract

Although the efficacy of vaccines targeting SARS-CoV-2 is apparent now that the approved mRNA and adenovirus vector vaccines are in widespread use, the longevity of the protective immune response and its efficacy against emerging variants remains to be determined. We have therefore designed a DNA vaccine encoding both the SARS-CoV-2 spike receptor-binding domain (‘RBD’) and nucleocapsid proteins, the latter of which is highly conserved amongst beta coronaviruses. The vaccine elicits strong pro-inflammatory CD4^+^ Th1 and CD8^+^ T-cell responses to both proteins in mice and rats, with responses being significantly enhanced by fusing the nucleocapsid sequence to a modified Fc domain. We have shown that the vaccine also stimulates high titre antibody responses to RBD in mice that efficiently neutralise in pseudotype and live virus neutralisation assays and show cross reactivity with spike proteins from the variants B.1.1.7 (Alpha), B.1.351 (Beta) and B.1.617.2 (Delta). The vaccine also showed good protection in a viral challenge model in ACE2 receptor transgenic mice. This DNA platform can be easily adapted to target variant proteins and we show that a vaccine variant encoding the Beta variant sequence stimulates cross-reactive humoral and T cell responses. These data support the translation of this DNA vaccine platform into the clinic, thereby offering a particular advantage for rapidly targeting emerging SARS-CoV-2 variants.

## Introduction

The SARS-CoV-2 pandemic has prompted a global response, with researchers from hundreds of institutions across the world developing and testing potential vaccine candidates that can prevent infection and coronavirus disease 2019 (COVID-19). SARS-CoV-2 is known to facilitate host cell entry by targeting the cellular receptor angiotensin-converting enzyme 2 (ACE2) using its densely glycosylated spike (S) protein. The S protein is a trimeric class I fusion protein that undergoes several structural rearrangements and cleavage events to eventually fuse the viral and host membranes ^1,2^, a process which is triggered by S1 subunit binding to the host cell receptor ACE2 via its receptor binding domain (RBD). The virus targets the airway epithelial cells, alveolar epithelial cells, vascular endothelial cells and macrophages in the lung, all of which express ACE2 ^3,4^. Viral assembly inside the host cell requires four structural proteins, the S, matrix (M), envelope (E), and nucleocapsid (N) proteins. Homotrimers of the S protein make up the receptor-binding spikes on the viral surface ^5,6^. The N protein contains two domains that each bind the viral genome, packaging it into virions ^7–9^. SARS-CoV-2 has a single-stranded positive sense RNA genome and the coronavirus families have the largest genomes in this category of viruses, encoding RNA processing and editing enzymes that increase the fidelity of replication. However, variation arises through a high rate of homologous recombination. Although SARS-CoV-2 can cause severe symptoms and death in the elderly or clinically vulnerable populations, the fact that approximately 80% of infected individuals present with no symptoms or mild infection ^10^ suggests that effective immunity can control the infection.

As the S protein has been shown to be the primary target for neutralising antibodies in patients with SARS-CoV-2 ^11^, it has been the focus of most vaccine development programmes. Although, the S protein is also a key target for T cell responses, responses have also been detected against the M, E and N proteins and several other non-structural proteins ^12^. The currently approved SARS-CoV-2 vaccines and most of those under development predominantly aim to stimulate neutralising antibody responses against the S protein to block the virus binding to ACE2 receptor and entering cells. However, viral evolution has created numerous mutations in the S protein that not only have increased infectivity but have also reduced the efficacy of neutralising antibodies that have been raised to previously encountered viruses ^13–16^. The N protein plays a vital role in viral RNA replication and the formation of new virions and is therefore highly conserved between coronaviruses ^4,17^. It is present in large quantities and can stimulate a strong immune response. Critically, most survivors of SARS-CoV-2 show evidence of T cell and antibody immunity against the N protein ^18^, suggesting that it is a target of the natural immune response. Our vaccine design includes the N protein in order to provide broader immunity, a strategy which is also being considered by others developing ‘next generation’ vaccines ^19^.

We have previously shown that delivery of epitopes from tumour antigens, linked to human immunoglobulin G1 (IgG1) fragment crystallisable (Fc) in a DNA vector, stimulates high avidity CD8^+^ T cell responses that are efficiently recruited into the memory pool ^20–22^. The DNA vector enables direct presentation of the encoded epitopes in antigen presenting cells (APCs) as well as cross presentation of secreted Fc-fused proteins to dendritic cells (DCs) via high affinity FcγR1 receptor (CD64) binding. This DNA vaccine platform has been successfully used in the clinic as a cancer immunotherapeutic, where it has been shown to induce vaccine-specific T cell responses in patients with melanoma and to be compatible with a prime and multiple boosts dosing regimen ^23^. Additionally, we have recently described an Fc-engineering approach that increases the functional avidity of antibody target binding by enhancing Fc:Fc co-operativity ^24^, and this modified Fc forms part of our vaccine design. Herein, we describe the validation of this versatile and rapid to manufacture DNA platform for generating a SARS-CoV-2 vaccine targeting both the N protein and the S RBD. We report that the linkage of the N protein to the modified Fc enhances T cell immunity and show that the inclusion of this N-modified Fc fusion alongside the RBD antigen in a bivalent DNA vector stimulates strong cellular and humoral immunity to both antigens. High titre antibody responses that exhibited pseudotype neutralisation with a 50% reduction in infective dose (ID_50_) of >5000 as well as protection against infection *in vivo* were induced and high frequency CD8^+^ and Th1 CD4^+^ T cell responses to both RBD and N proteins stimulated. Finally, we demonstrate the induction of variant cross-reactive immune responses and the flexibility of this DNA platform for targeting SARS-CoV-2 variants. Collectively, these results underpin the advancement of this DNA vector encoding RBD and modified Fc-linked N protein into clinical trials.

## Results

### Design and characterisation of DNA constructs

Spike RBD and N protein DNA sequences were obtained from the Lineage A SARS-CoV-2 variant and engineered alongside N terminal human IgH leader sequences into the pVAX1 vector backbone. Short (aa330-525) or long (aa319-541) S protein RBD sequences were fused in frame with human IgG1 Fc constant region or a fibritin timer domain or disulphide bridge motif (Figure 1Ai and Supplementary Figure 1A). N protein sequences were added alongside the S protein RBD sequences, alone or fused in frame with human IgG1 Fc (Fc) constant region or a modified human IgG1 Fc (Fc iV1) constant region (Figure 1Ai). A commercially available vector encoding the whole S sequence was also used (Figure 1Aii).

**Figure 1.**
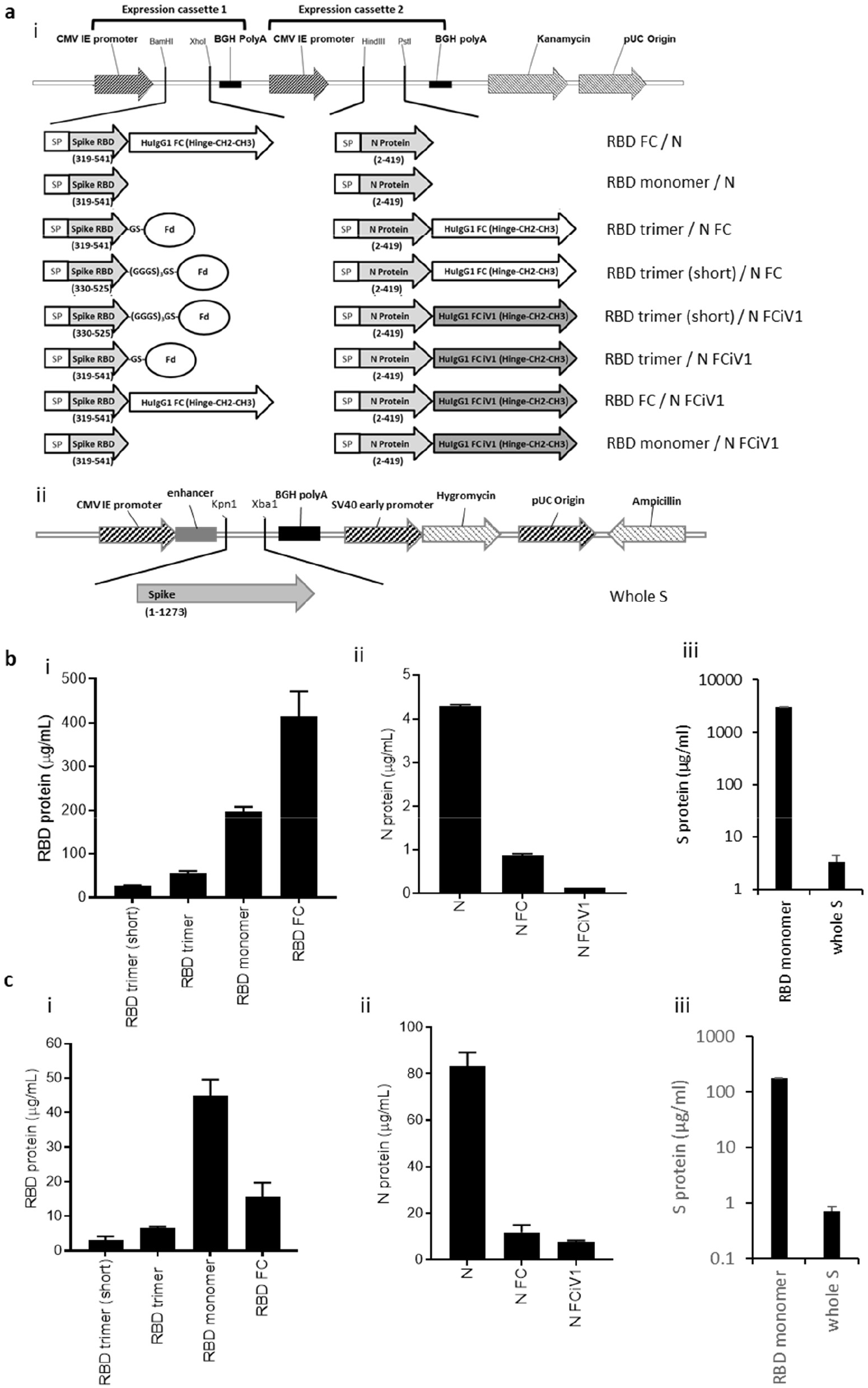
Schematic representation of the DNA SARS-CoV-2 plasmids and expression of N and RBD proteins. A, DNA constructs expressing N and RBD variants (i) and whole S (ii). Spike RBD and NP variant chains are within expression cassettes 1 and 2 of the pVaxDC vector. Numbers indicate amino acids. SP: signal peptide/ Human IgH Leader, RBD: receptor-binding domain, Fd: Fibritin fold on trimer motif from T4 bacteriophage (GYIPEAPRDGQAYVRKDGEWVLLSTFL) is attached to the S RBD via glycine/serine linkers. In some constructs the Spike RBD domain and N protein is fused inframe with HuigG1 FC or the improved modified version of the constant domain, designated iV1. RBD (i) and N (ii) protein expression from vaccine constructs or S protein expression from whole S DNA (iii) in transfected Expi293™ cells in the supernatant (B) and cell lysate (C). Data are representative of at least 2 independent experiments.

Expression of both N protein and RBD or S protein from DNA constructs was assessed by transfection of Expi293F™ cells followed by analysis of supernatant and cell lysate samples by sandwich ELISA. Constructs expressing the RBD region as a monomer or linked to Fc showed the highest levels of RBD protein secretion by Expi293F™ cells, whereas those containing the long RBD trimer or shorter RBD trimer versions exhibited lower protein secretion levels (Figure 1Bi). No significant difference was seen between the trimer variant constructs (Supplementary Figure 1B). Overall, the levels of N protein secretion were much lower than those for RBD, with the non-fused N protein producing more secreted protein than the Fc and Fc iV1 N protein constructs (Figure 1Bii). The RBD monomer was secreted at higher levels to the whole S protein (Figure 1Biii). Unlike antibody responses, the stimulation of a T cell response is not reliant on secreted protein. To determine if protein was made but not secreted we therefore also measured expression levels in cell lysates. Cells transfected with the construct expressing the RBD monomer showed the highest level of RBD protein in the cell lysates (Figure 1Ci). A similar observation was seen for the N protein, with highest levels in the cell lysate transfected with the construct expressing the non-fused N protein (Figure 1Cii). Similarly, analysis of S protein in the cell lysates from cells transfected with RBD monomer or whole S DNA showed higher protein expression from the RBD monomer (Figure 1Ciii). Collectively, the results suggest that functional RBD and N proteins are produced as both secreted and intracellular forms, which could stimulate both T cell and antibody responses.

### Linkage of N protein to modified Fc stimulates superior N specific T cell responses

T cell responses in HLA-A2 transgenic mice following immunisations (schematic shown in Figure 2A) with vaccine constructs containing the N monomer, monomer linked to Fc or modified Fc (iV1) were evaluated using RBD and N protein overlapping peptide pools. Peptide pools contain 15aa peptides that overlap by 11aa and span the entire sequence and stimulate both CD4^+^ and CD8^+^ T cell responses without requiring any prior knowledge of the presenting MHC allele. Splenocytes were isolated from mice 6 days after the last immunisation and T cell responses following stimulation assessed *ex vivo* using an IFNγ ELISpot. In addition to the peptide pools, several peptides were selected from published reports and using prediction algorithms that were known, or suggested to bind to HLA-A2 or murine H-2Kb/d, H-2Db/d, I-Ab or I-Ad/Ed (Table S1).

**Figure 2.**
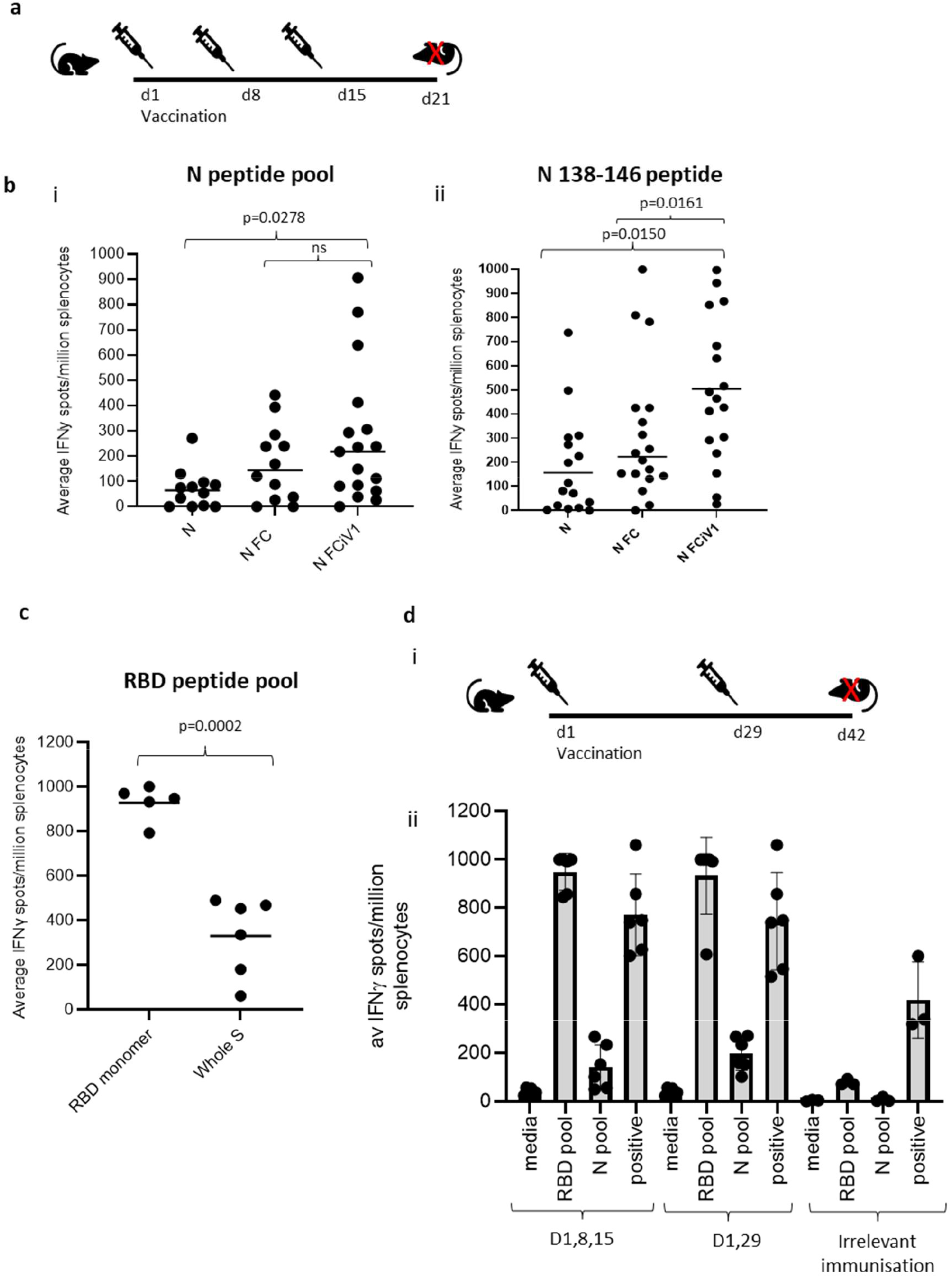
Linkage of N protein to modified Fc stimulates superior N and RBD specific T cell responses. A, Schematic of the immunisation regime. B, BALB/c, C57Bl/6 or HLA-A2 transgenic mice immunised with DNA constructs expressing N, N linked to Fc or N linked to modified Fc (iV1) via gene gun and T cell responses monitored by IFNγ ELISpot assay to N peptide pool (i) or N 138-146 peptide (ii). C, Assessment of RBD specific T cell responses using RBD peptide pool from constructs containing RBD monomer alongside N linked to modified Fc (iV1) compared to DNA encoding the whole S sequence. Responses are measured as spots/10^6^ splenocytes and normalised against unstimulated controls for comparison. D, Comparison of an alternate immunisation regime (i) for IFNγ responses to RBD and N peptide pools compared in BALB/c mice immunised with DNA construct expressing RBD monomer and N linked to modified Fc (iV1) via gene gun or irrelevant DNA immunisation (ii). Symbols represent mean response for individual mice, line/bar represents mean value between mice. Significant P values are shown from studies with matched data points. Data are collated from independent studies, in which n=3.

A high frequency of T cells specific to the N protein was detected in mice immunised with DNA constructs containing the N protein linked to modified Fc (iV1) when compared to constructs containing non-fused N protein (*p*=0.0278) or the N protein linked to unmodified Fc, although the latter did not reach significance (Figure 2Bi). A high frequency of T cells recognising the N 138-146 peptide was induced in HLA-A2 transgenic mice following immunisation with the DNA construct containing the N protein linked to a modified Fc (iV1) when compared to the frequency observed in mice immunised with the constructs containing non-fused N protein (*p*=0.0150) or linked to Fc (*p*=0.0161) (Figure 2Bii). The peptide (N 138-146), which is conserved between SARS-CoV and SARS-CoV-2 ^25^, stimulated a response in cells from immunised HLA-A2 transgenic mice, thereby highlighting the potential of these T cells to recognise other coronaviruses.

A high frequency of T cells specific to the RBD antigen was detected in mice immunised with the vaccine constructs. The construct containing the RBD monomer induced a higher frequency of T cells specific for the RBD protein when compared to the frequency of responses induced by a DNA construct containing the whole S sequence (*p*=0.0002) (Figure 2C), possibly due to either focussing of the immune response on the RBD region rather than other epitopes outside the RBD region or due to differences in the level of protein expression. In addition, T cell responses induced by the RBD monomer were similar to those seen from constructs containing the RBD trimer variants and did not appear to be associated with lower levels of protein expression (Supplementary Figure 1B and C).

To examine if any difference in T cell responses is observed with longer immunisation regimes, responses were compared in BALB/c mice after immunisation with the construct containing the RBD monomer and N linked to modified Fc (iV1). Immunising at days 1, 8 and 15 was compared to two immunisations at days 1 and 29 (Figure 2Di). No differences in levels of T cell responses to either RBD or N peptide pools were seen (Figure 2Dii). In addition, minimal responses were seen in mice immunised with an irrelevant DNA construct. To confirm that T cell responses could also be stimulated in different species, responses were also analysed in immunised rats (Supplementary Figure 2A). These results demonstrated that T cell responses to RBD and N protein could be induced in rats.

T cell responses were further evaluated in BALB/c, C57Bl/6 and HLA-A2 transgenic mice following immunisation with the DNA construct expressing RBD alongside N linked to modified Fc (iV1) (also known as SN15) using RBD and N peptide pools and individual peptides. A high frequency of T cells specific for SARS-CoV-2 N and RBD were detected across all strains of mice. T cell responses to two longer peptides (RBD 505–524 and N 80–100) in addition to the RBD and N peptide pools were seen in BALB/c mice (*p*=0.0012, *p*=0.0171, *p*=0.0001 and *p*=0.0368, respectively) (Figure 3A). Immunised C57Bl/6 mice also exhibited T cell responses to the RBD 505–524 peptide and lower frequency responses, which did not reach statistical significance, to the N 212–231 and N 101–121 peptides (Figure 3B). T cell responses to the two short peptides RBD 417-425 (*p*<0.0001) and N 138-146 (*p*<0.0001) peptides were detected in immunised HLA-A2 transgenic mice (Figure 3C) - these peptides have been described as potential HLA-A2 epitopes in human studies ^25,26^. To confirm these to be CD8-binding peptides, the T cell responses were assessed in the presence of CD4 or CD8 blocking antibodies (Figure 3D). The presence of the CD8 blocking antibody totally abrogated both responses (*p*<0.0001), whereas the CD4 blocking antibody had little effect, thus confirming that these were CD8^+^ T cell responses in HLA-A2 transgenic mice.

**Figure 3.**
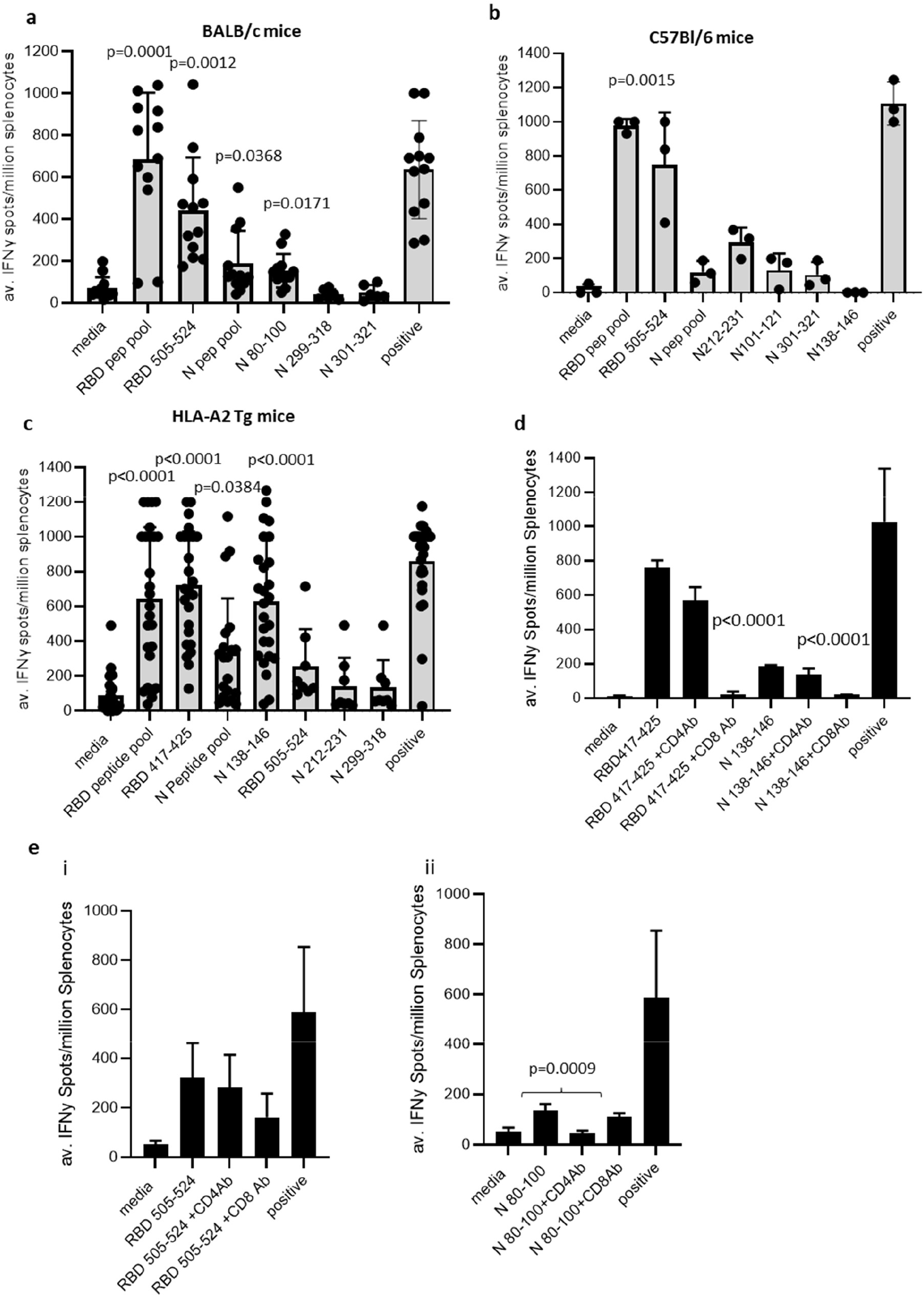
DNA vaccination induces strong CD8^+^ and CD4^+^ T cell responses. BALB/c (A), C57Bl/6 (B) or HLA-A2 transgenic (C) mice were immunised on days 1, 8 and 15 with DNA constructs expressing RBD alongside N linked to modified Fc (iV1) using a gene gun and T cell responses to RBD or N peptide pools and individual peptides monitored by IFNγ ELISpot assay on day 21. Symbols represent mean response for individual mice, line represents mean value between mice. Data are collated from multiple independent studies and p values shown are compared to medium control. D, Responses from HLA-A2 transgenic mice to RBD 417-425 or N 138-146 peptides assessed in the presence of CD4 or CD8 blocking antibodies. E, Responses from BALB/c mice to RBD 505-524 (i) or N 80-100 (ii) peptides assessed in the presence of CD4 or CD8 blocking antibodies. Results are representative of at least two independent experiments in which n=3 mice per group. Responses are measured as spots/10^6^ splenocytes.

To characterise the T cell responses to the longer peptides which have the potential to contain both CD8 and CD4 epitopes, BALB/c mice were immunised with DNA constructs expressing RBD alongside N linked to modified Fc (iV1) and T cell responses to the N 80-100 and RBD 505-524 peptides determined by IFNγ ELISpot assay in the presence of CD4 or CD8 blocking antibodies. A strong T cell response to the N 80-100 peptide was observed and this was efficiently blocked by the CD4 antibody (*p*=0.0009), but not the CD8 antibody, implying the presence of a CD4 epitope within this peptide sequence (Figure 3Eii). Although a strong T cell response to the RBD 505–524 peptide was observed, this was not significantly inhibited by either the CD4 or CD8 blocking antibodies, suggesting that this peptide sequence stimulates both CD4 and CD8 responses (Figure 3Ei).

To confirm both CD4^+^ and CD8^+^ T cell responses to the RBD and N proteins, splenocytes from BALB/c and C57Bl/6 mice immunised with the DNA construct expressing RBD alongside N linked to modified Fc (iV1) were expanded *ex vivo* and responses measured by intracellular IFNγ staining. Populations of CD4^+^ and CD8^+^ T cells produced IFNγ in response to restimulation with the RBD or N peptide pools as well as to the individual RBD 505-524 and N 80-100 peptides (Figure 4A, Supplementary Figure 3), thus confirming the presence of both CD4^+^ and CD8^+^ T cell responses to both antigens. To confirm the polarisation of the responses, analysis of other cytokines *ex vivo* using the Mesoscale Discovery™ platform was performed. Analysis of supernatant from the IFNγ ELISpot assay plates showed that in addition to IFNγ, IL-10 and TNFα were detected after stimulation with the RBD peptide pool, as well as lower levels of IL-1β, IL-17A and IL-2 (Figure 4B). Stimulation with the N peptide pool showed low levels of IL-10 and IL-2 in addition to IFNγ. No IL-4 was detected in response to either RBD or N peptide pool stimulation. A similar cytokine profile was also shown by intracellular cytokine staining (Supplementary Figure 3). This data provides evidence in two different mouse models that CD4^+^ and CD8^+^ T cell responses can be stimulated by vaccination to both the N and RBD proteins and that CD4^+^ T cell responses are skewed toward Th1 rather than Th2.

**Figure 4.**
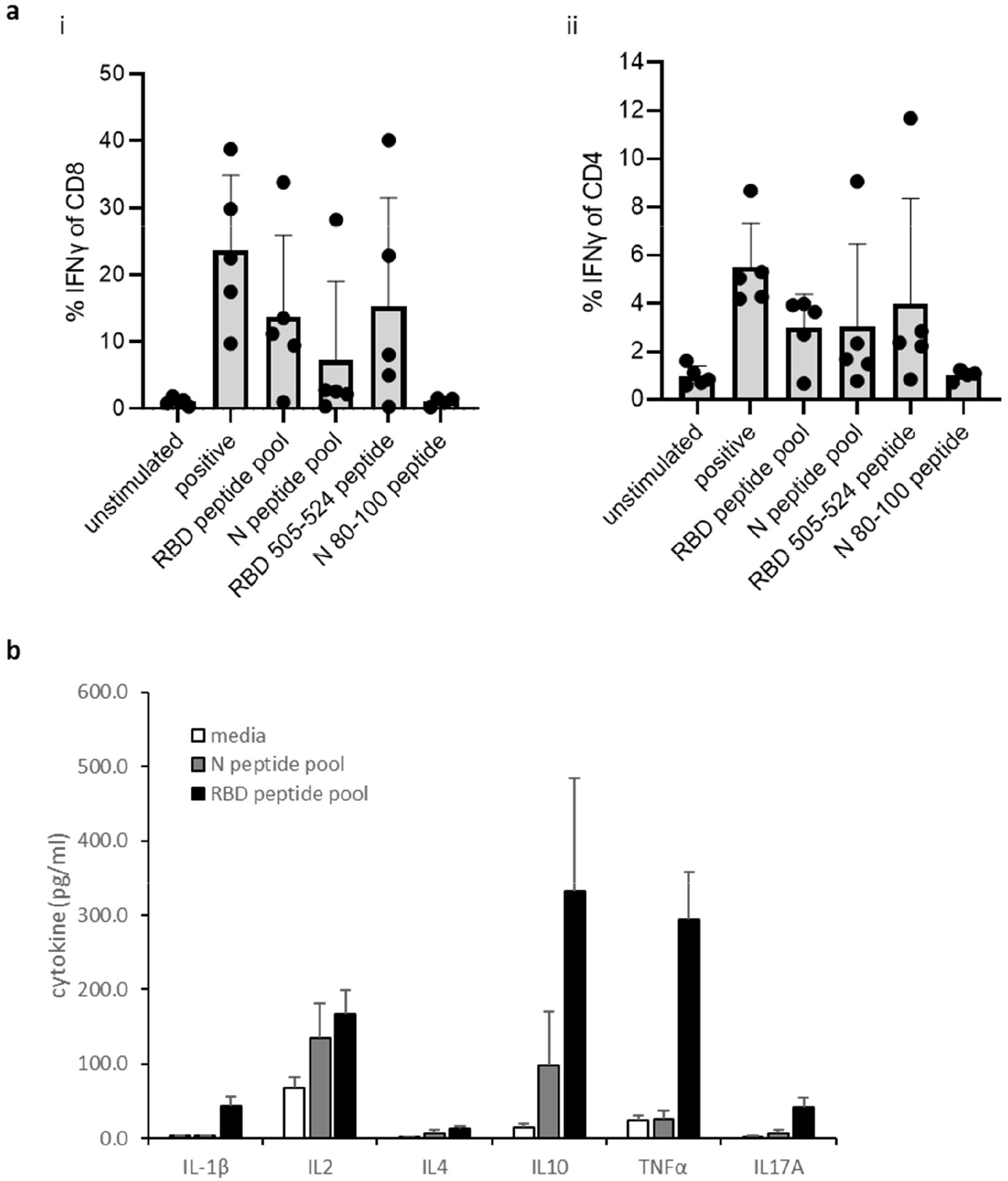
CD4^+^ and CD8^+^ T cell responses with Th1 polarisation. A, Intracellular cytokine staining for IFNγ, in response to peptide pool or individual peptide restimulation or stimulation with anti-CD3 antibody positive control on splenocyte cultures expanded *in vitro* with S1 and N proteins and peptide pools from BALB/c and C57Bl/6 mice immunised with DNA construct expressing RBD alongside N linked to modified Fc (iV1). Percentages of CD4^+^ or CD8^+^ cells T cells are shown as averages from five independent experiments, for each of which n=3. B, cytokines detected using the Mesoscale Discovery platform in culture supernatants after *ex vivo* stimulation on day 21 of splenocytes from BALB/c mice immunised on days 1, 8 and 15 with DNA construct expressing RBD alongside N linked to modified Fc (iV1) with an RBD and N peptide pool, n=6.

### DNA vaccination stimulates strong N and RBD binding antibody responses that show efficient pseudotype neutralisation and protection against viral challenge

Humoral responses were evaluated in sera from BALB/c mice immunised three times with constructs containing the RBD monomer, whole S DNA, RBD trimer or RBD monomer linked to Fc and analysed for reactivity to SARS-CoV-2 S1 protein by ELISA (Figure 5Ai and ii). Vaccination induced potent antibody responses to the S1 protein that were detectable in sera against all constructs up to a 1 in 100,000 dilution. Constructs containing the RBD monomer and RBD linked to Fc showed the highest EC_50_ and quantity of immunoglobulin (Ig), with 58.2-59.9μg/mL anti-S1 Ig being detected. Sera were also analysed for reactivity to SARS-CoV-2 N protein by ELISA. Lower antibody responses were detected against the N protein compared to RBD, with similar EC_50_ values obtained for all constructs irrespective of whether the N protein was fused to Fc or not (Figure 5B).

**Figure 5.**
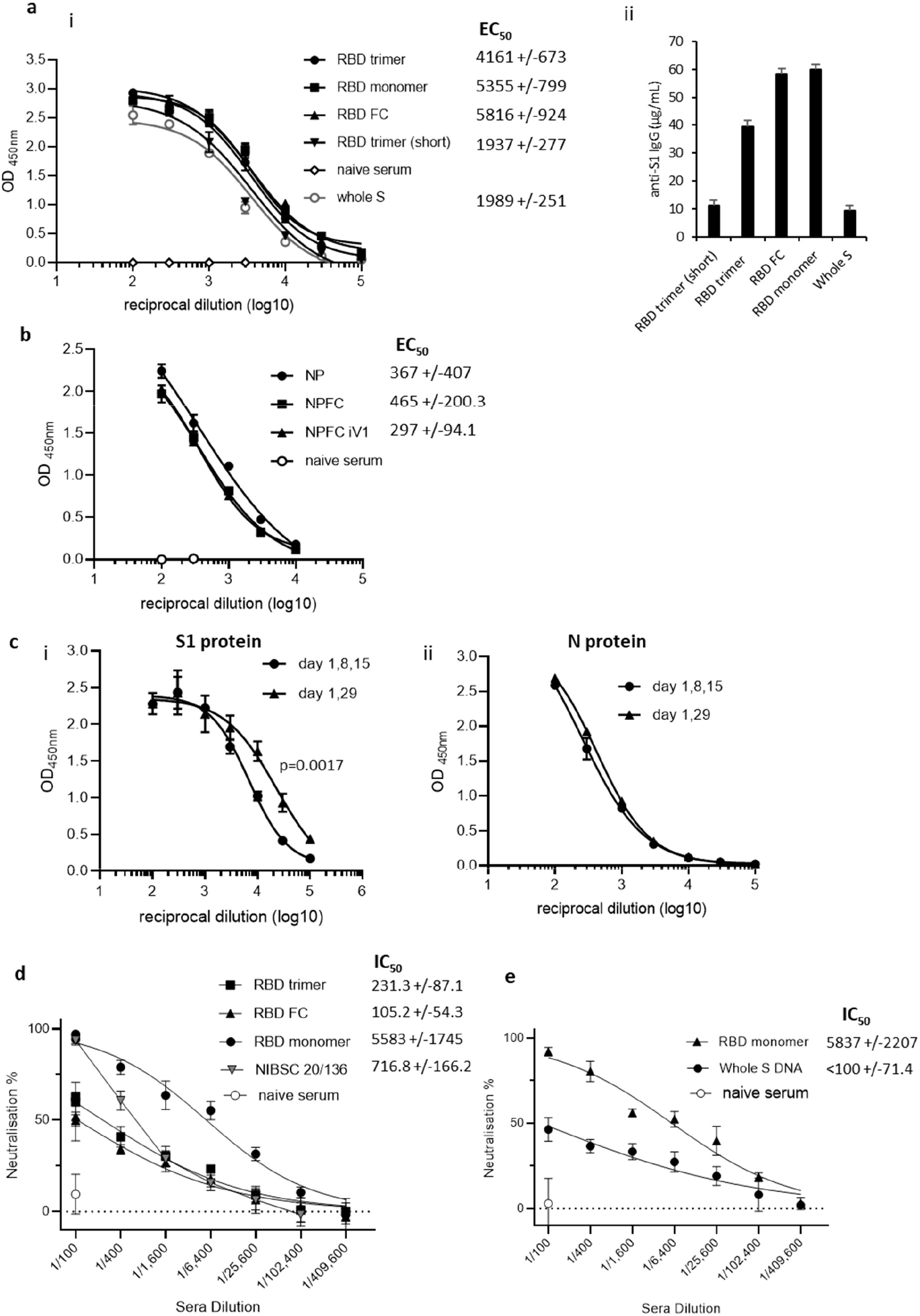
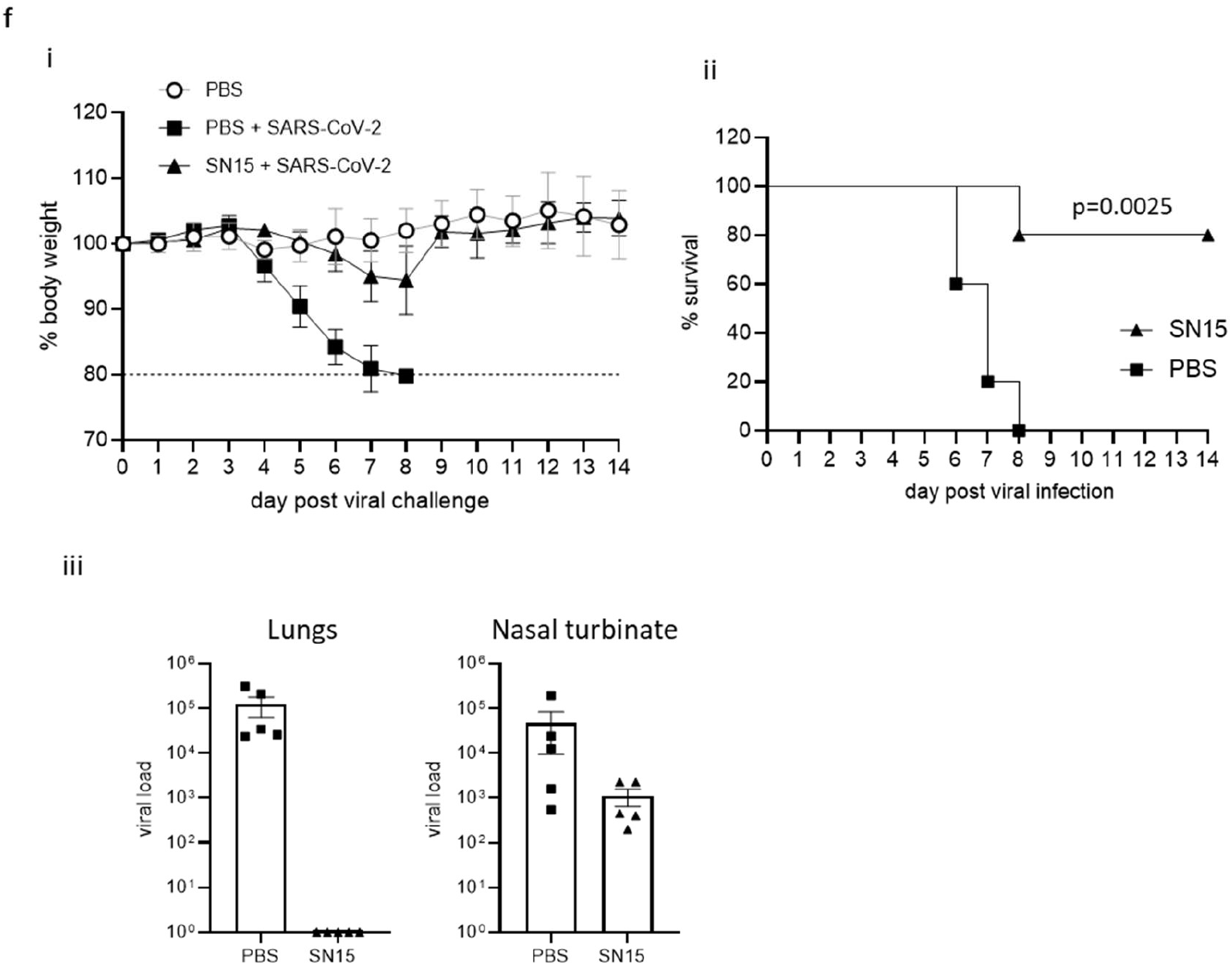
DNA vaccination induces anti-RBD neutralising and anti-N antibody responses. Sera from mice immunised on days 1, 8 and 15 using a gene gun analysed for anti-S1 (A) or anti-N (B) antibodies by ELISA on day 21. Data presented show absorbance readings with sera titrations (i) and as total μg/mL specific IgG (ii) EC_50_ values were calculated and shown with 95% Confidence Interval as a measure of variation. The quantity of immunoglobulin (Ig) in the sera was estimated from the OD value at a 1 in 3000 serum dilution using the standard curve of commercial murine S1 antibody. C, Comparison of an alternate immunisation regime of day 1 and 29 using a construct expressing RBD monomer and N linked to modified Fc for antibody responses to (i) S1 and (ii) N protein. Sera from BALB/c mice immunised with constructs containing the RBD monomer, trimer or monomer linked to Fc (D), C57Bl/6 mice immunised with construct containing the RBD monomer or whole S protein (E) and the NIBSC 20/136 standard were assessed for pseudotype neutralisation at increasing sera dilutions and IC50 values displayed. Data are representative of at least two independent studies, for each of which n=3. F, K18 hACE2 transgenic mice were challenged with Lineage A virus strain after vaccination with DNA construct expressing original variant RBD alongside N linked to modified Fc (iV1) (SN15) and bodyweights (i), survival (ii) and viral load (iii) monitored, N=5.

To examine if any difference in antibody responses is observed with a longer immunisation regime, responses were compared in BALB/c mice after immunisation with three doses of the construct containing the RBD monomer and N linked to modified Fc (iV1). For this, immunising on days 1, 8 and 15 was compared to immunising on days 1 and 29. Although increased titres of S1 specific antibody responses were observed using the day 1 and 29 immunisation regime, antibody responses to N protein remained similar (Figure 5C). A longer-term analysis of S1 specific antibody responses reveals a small decrease by day 82 after immunisations at days 1 and 29, but this can be efficiently increased with a booster immunisation at day 85, whereas antibody responses to N protein remained unchanged (Supplementary Figure 4). We also show that S1 and N specific antibody responses can be induced in rats (Supplementary Figure 2B).

The generation of a potent antibody response which is capable of inhibiting the binding of the S protein to ACE2 receptor and subsequent viral infection is vital for an effective SARS-CoV-2 vaccine. The ability of sera from mice immunised with the DNA constructs containing RBD monomer, trimer or monomer linked to Fc to prevent viral infection was therefore tested in a pseudotype neutralisation assay. Sera from mice immunised with the RBD monomer showed higher neutralisation IC_50_ values (Figure 5D), with 50% neutralisation at serum dilutions of greater than 1 in 5000 - higher than that seen with the NIBSC 20/136 international antibody standard. The sera showed no neutralisation of an irrelevant pseudotype virus and no neutralisation was seen using serum from naïve mice (Supplementary Figure 5). Sera from mice immunised with the RBD monomer DNA construct showed higher IC_50_ neutralisation values compared to that from mice immunised with the DNA construct expressing the whole S (Figure 5E). This is also evident from the differing antibody levels shown in Figure 5Aii where the RBD monomer generated high EC_50_ values when compared to the construct expressing full-length S protein.

The construct encoding the RBD monomer alongside the N linked to modified Fc (iV1) (SN15) was tested for efficacy against SARS-CoV-2 in transgenic mice expressing human angiotensin-converting enzyme 2 (hACE2) driven by the human cytokeratin 18 promoter (K18 hACE2). Upon viral challenge, control (PBS treated) mice challenged with virus show rapid loss of body weight (Figure 5Fi) whereas SN15 DNA vaccinated mice maintain body weights similar to unchallenged controls. This protection against viral challenge is also demonstrated by significantly enhanced survival (Figure 5Fii) and total elimination of viral load in lungs as well as reduction of viral load in nasal turbinate, at day 2 post infection (Figure 5Fiii), in the vaccinated group compared to challenged control providing evidence that the DNA vaccination prevents infection and disease *in vivo*.

### Responses induced by DNA vaccination show variant cross reactivity

The emergence of new SARS-CoV-2 variants is a major cause for concern as they have the potential for greater transmissibility and increased disease severity ^15,27,28^. Variants of SARS-CoV-2 have the potential to impact the efficacy of the current vaccines targeting the original Lineage A virus and although protection against hospitalisation is low, there remains some uncertainty about the effectiveness against these new variants ^29–31^. We therefore examined the ability of sera from BALB/c mice immunised with the DNA vaccine expressing RBD alongside N linked to modified Fc (iV1) (SN15) for reactivity to the Lineage A SARS-CoV-2 S1 protein as well S1 proteins incorporating the mutations seen in the Alpha, Beta and Delta variants by ELISA (described in Supplementary Table 3). Vaccination induced a potent antibody response with reactivity to both Alpha and Beta S1 proteins (Figure 6A) and Beta and Delta RBD proteins (Figure 6C) detectable in sera at up to a 1 in 100,000 dilution. However, in line with other reports, we also observed a significantly (*p*<0.0001) lower reactivity to the Beta variant S1 protein compared to the Lineage A protein, whereas reactivity to the Alpha variant S1 protein was not significantly lower. Reactivity to the Delta variant RBD protein was higher than that to the Beta variant RBD protein.

**Figure 6.**
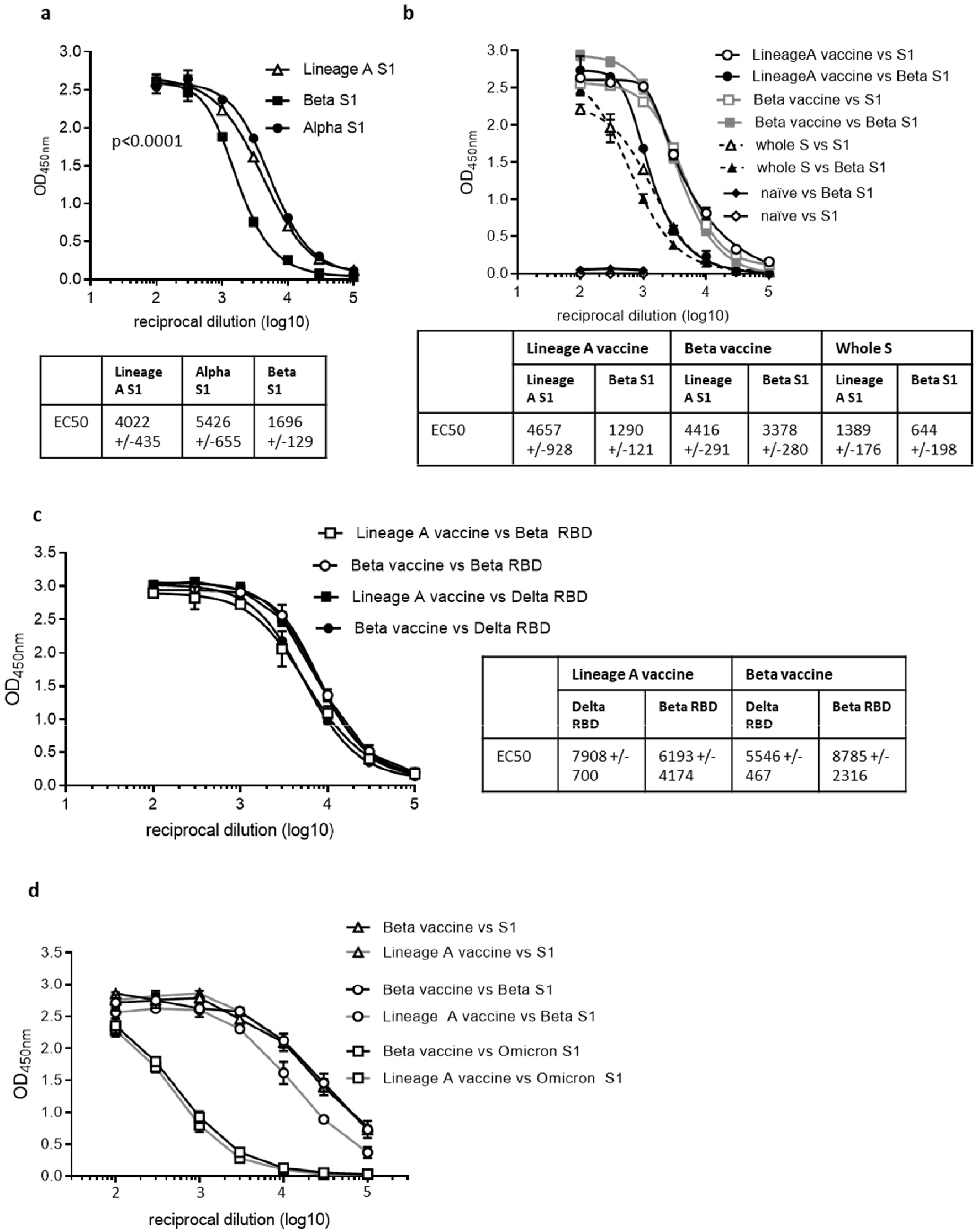
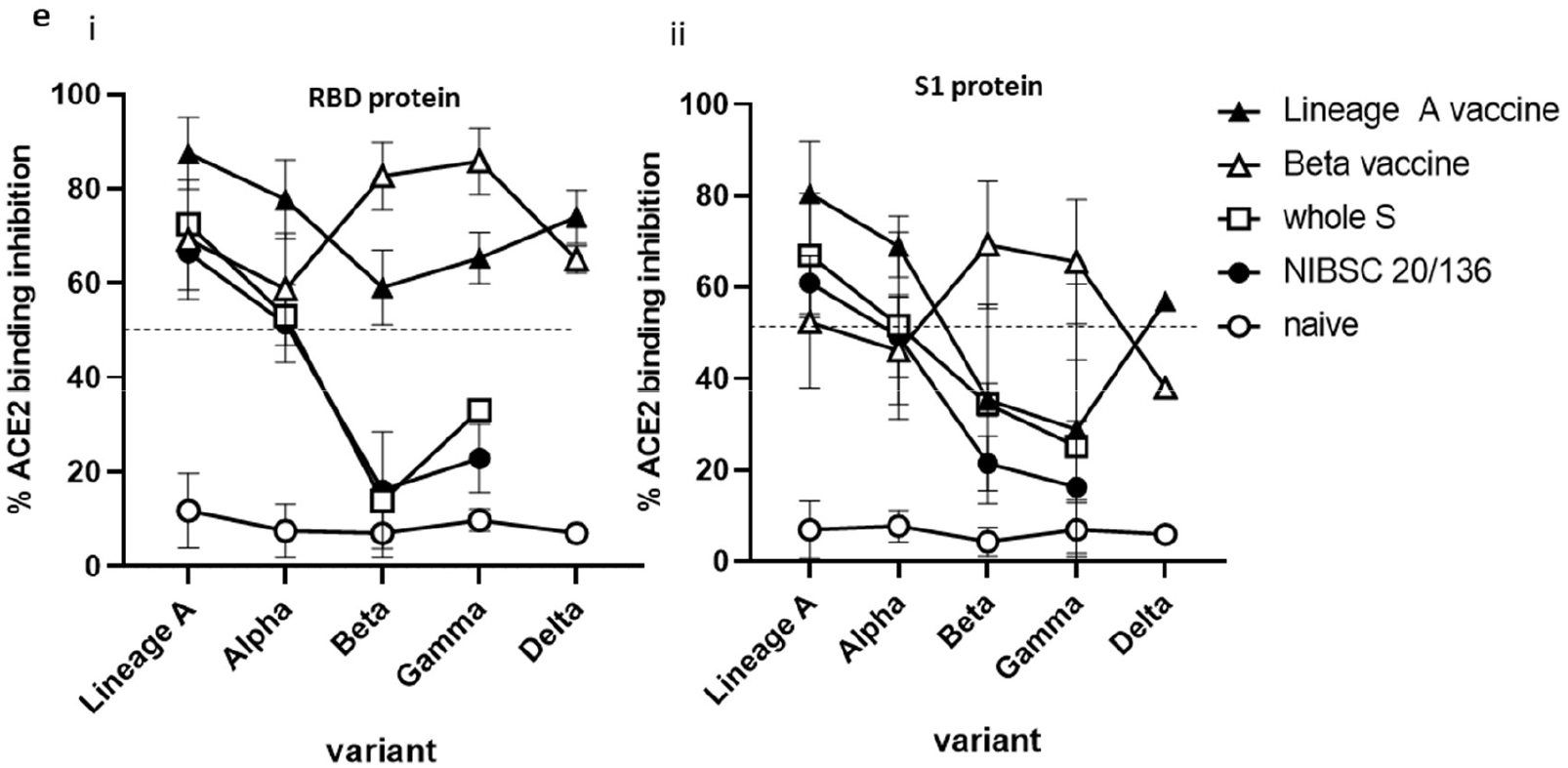
DNA vaccination induces cross reactive antibody responses. BALB/c mice were immunised on days 1, 8 and 15 with DNA constructs expressing original or Beta variant RBD alongside N linked to modified Fc (iV1) or whole S DNA (B and D) using a gene gun. Serum from immunised mice at day 21 were analysed for antibodies by ELISA at various sera dilutions using variant S1 proteins (A and B) or variant RBD proteins (C) and EC_50_ values were calculated and shown with 95% Confidence Interval as a measure of variation. Data are representative of at least two independent studies, for each of which n=3. Serum at day 82 from mice immunised at days 1 and 29 were analysed in ELISA at various sera dilutions using variant S1 proteins (D). Serum at days 21 or 42 from mice immunised at days 1, 8 and 15 or 1 and 29 respectively were analysed in ACE2 binding inhibition assay *versus* RBD or S1 protein at 1:100 sera dilution (Meso Scale Discovery Platform) (E). Data are collated from at least two independent studies, for each of which n=3.

Data have suggested that although the approved vaccines induce responses that can provide some cross-reactivity with the Alpha variant, they are less effective against the Beta, Delta and Gamma variants ^27,32–34^. We engineered a construct encoding the RBD containing the mutations within the Beta variant alongside N linked to modified Fc (iV1) (also known as SN17). Sera from Balb/c mice immunised with the Beta vaccine construct or a construct expressing the whole S protein from Lineage A were analysed for reactivity to the original Lineage A SARS-CoV-2 S1 protein as well S1 protein incorporating the mutations seen in the Beta variant or RBD protein from Beta and Delta variants by ELISA. High antibody responses were observed against the protein variants up to a 1 in 100,000 dilution (Figure 6B and C). The Beta variant vaccine generated higher titres to the B.1.351 (Beta) variant S1 protein (EC_50_ of 3931), with lower titres against the original Lineage A S1 protein (EC_50_ of 2086) (Figure 6B). Both vaccines induced higher titres to the original Lineage A S1 and Beta S1 proteins compared to responses induced by the whole S DNA vaccine (EC_50_ of 1389 and 644 respectively). Reactivity to the Delta variant RBD protein was assessed by ELISA and compared to reactivity to the Beta variant RBD protein. Figure 6C demonstrates stronger antibody reactivity to the Beta RBD protein compared to the Delta RBD protein. Despite these differences, sera from mice immunised with either vaccine show high titre antibody responses to variant RBD proteins. Sera from mice vaccinated with the Lineage A and Beta vaccines were also assessed for reactivity to the Omicron variant (B.1.1.529) S1 protein and although endpoint titres of up to 1 in 10,000 are observed the reactivity demonstrates a drop in response by 1-2 log values compared to Lineage A or Beta S1 proteins (Figure 6D), an observation consistent with other vaccine reports ^35,36^.

The sera were also assessed in a similar antibody binding assay on the MesoScale Discovery platform that includes the whole spike and RBD proteins from original Lineage A, Beta, Alpha, Delta and P.1 (Gamma) variants (Supplementary Figure 6) where a similar pattern of response was observed. Sera from immunised mice showed similar reactivity to either the original Lineage A and Alpha variants or the Beta and Gamma variants, as might be expected based on their RBD mutations, however both vaccines demonstrate similar responses to the Delta variant proteins. Interestingly, the sera from mice vaccinated with the original Lineage A vaccine and the Beta variant vaccine showed similar titres of antibodies recognising the Lineage A N protein and a N protein variant containing the D3L, R203K, G204R and S235F mutations, two of which are also found in the Alpha variant (Supplementary Figure 7), suggesting perhaps that N protein specific antibodies may be more cross-reactive or bind to conserved regions of the N protein.

The ability of sera from immunised mice to inhibit binding of the variant RBD or whole S proteins to the ACE2 receptor was assessed using the MesoScale Discovery platform. Inhibition of RBD variant binding to ACE2 was higher for sera from mice vaccinated with the original Lineage A vaccine construct and the Beta variant vaccine construct compared to that seen with the NIBSC 20/136 control (Figure 6E). In contrast, sera from mice immunised with whole S DNA showed a lower capacity to inhibit ACE2 binding to variant proteins, which was similar to that of the NIBSC 20/136 control. Sera from mice immunised with the original Lineage A vaccine construct inhibited 80–100% of ACE2 binding to the original Lineage A and Alpha variant RBD, and inhibited binding to the Beta and Gamma RBD variants by 50–60% (Figure 6Ei). The reverse is seen in sera from mice immunised with the Beta variant vaccine. Despite a reduction in the inhibition of ACE2 binding to the Beta and Gamma RBD variants, the inhibition levels remain above those seen using sera from whole S DNA immunised mice. A similar trend is seen in the ACE2 receptor binding inhibition assay with the variant whole S proteins (Figure 6Eii).

Sera from mice immunised with the original Lineage A or Beta variant vaccines were also assessed in pseudotype and live virus neutralisation tests against the original Lineage A and Beta variants. Sera from mice immunised with the Lineage A variant vaccine showed potent neutralisation of the Lineage A pseudotype, with reduced efficacy against the Beta variant vaccine (ID_50_ values of 6232 and 2137, respectively) (Figure 7A). However, due to variation between replicates, this difference does not reach statistical significance. Sera from mice immunised with either vaccine showed neutralisation of the Beta pseudotype variant, but little difference was noted (ID_50_ values of 948 and 997, respectively) and no response was seen in serum from naïve mice.

**Figure 7.**
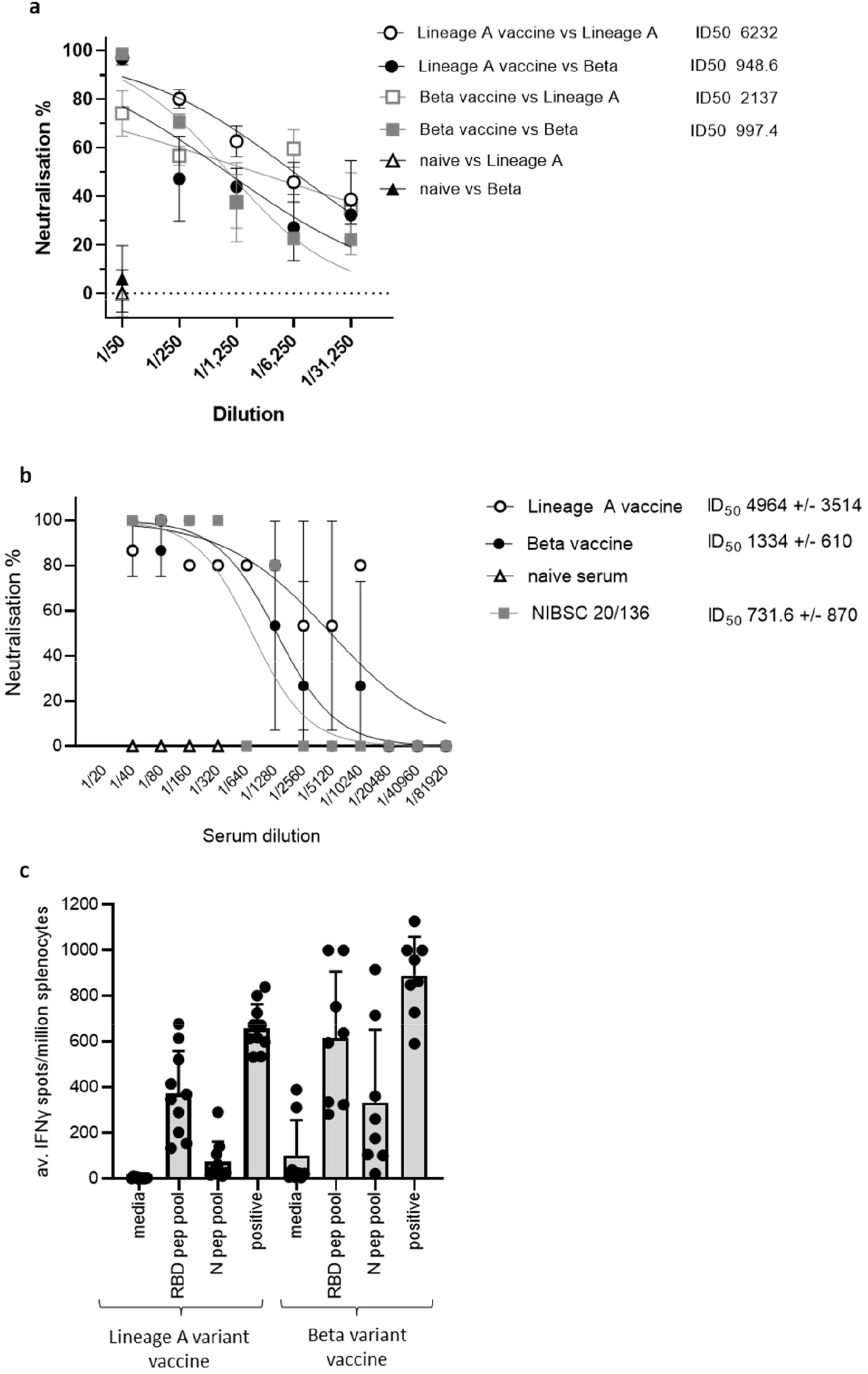
DNA vaccination induces cross-reactive neutralising antibody responses and cross-reactive T cell responses. BALB/c mice were immunised on days 1, 8 and 15 with DNA constructs expressing original or Beta variant RBD alongside N linked to modified Fc (iV1) using a gene gun. Sera from immunised mice at day 21 were analysed in pseudotype neutralisation assay against Lineage A and Beta pseudotypes (A) or a live virus neutralisation assay against Lineage A virus (B) and activity compared to naïve sera and NIBSC 20/136 reference standard. IC50 values were calculated and shown with 95% Confidence Interval as a measure of variation. Data are representative of at least two independent studies, for each of which n=3. C, T cell responses to RBD or N peptide pools in immunised mice monitored by IFNγ ELISpot assay at day 21. Symbols represent mean response for individual mice, line represents mean value between mice. Data are collated from at least two independent studies, for each of which n=3.

In a live virus neutralisation assay, sera from mice immunised with either the original Lineage A or the Beta variant vaccines neutralised the original Lineage A virus with ID_50_ values of 4964 and 1334 respectively, and both are higher than the National Institute for Biological Standards and Control (NIBSC) controls (Figure 7B). No neutralisation was seen using naïve mouse sera. This compares favourably with NIBSC 20/150 human convalescent plasma, which shows high neutralising antibody responses, and is higher than NIBSC 20/148, which shows moderate neutralising antibody responses (Supplementary Figure 8).

In contrast to the antibody responses, the T cell responses did not seem to be impacted by variations between the virus strains (Figure 7C). Splenocytes from mice immunised with either the original Lineage A vaccine or the Beta vaccine were stimulated *ex vivo* with RBD and N peptide pools derived from the original sequence. T cell responses specific for RBD and N were detected with little difference between the response frequency induced by the different vaccine constructs. This was also the case when responses were analysed over a range of antigen concentrations (Supplementary Figure 9). These results suggest that mutations in the Beta variant have less impact on the T cell responses.

## Discussion

We describe the development of a DNA vectored vaccine targeting both S and N proteins of SARS-CoV-2 and demonstrated that the vaccine induces CD4^+^ and CD8^+^ T cell responses to both N and RBD proteins, in addition to eliciting potent neutralising antibody responses. The stimulation of T cell responses in several mouse strains, as well as in rats, indicates the application of this vaccine in several species. Upon stimulation *ex vivo* T cells released IFNγ and TNFα, two Th1 cytokines required to induce potent immune responses, in the absence of IL-4 and potentially damaging Th2 responses. The induction of strong T cell immune responses by both Lineage A and Beta vaccines suggests that the mutations may not be as significant for evading T cell responses as they are for antibody responses. These data support the report from Tarke *et al*. suggesting SARS-CoV-2 variants have little impact on the T cell responses induced ^37^. A strong T cell response is not only essential for the generation of humoral immunity in coronavirus infection ^38^, but has also been shown in MERS-CoV and SARS-CoV infections to persist over time and is therefore likely to be important for long term protection ^18,39^. Grifoni *et al*. have indeed shown the importance of both T cell and humoral immunity to S and N proteins for SARS-CoV-2 protection ^12^. Therefore, to be most effective, immunity stimulated by vaccination will likely need to involve potent virus specific CD8^+^ and CD4^+^ T cell responses as well as neutralising antibody responses. Several reports using murine models indicate that in addition to CD8^+^ T cell responses, a strong Th1 response and virus specific neutralising antibodies are necessary for successful control of SARS-CoV and MERS-CoV ^40–42^. In humans, infection-specific memory T cells have been detected in convalescent patients and responses can be detected up to 18 weeks after infection. Responses to S protein in convalescent serum appear superior to those detected in individuals vaccinated with the currently approved vaccines ^43^, which is consistent with our own unpublished data. Studies by Ferretti *et al*. and Peng *et al*. reveal that SARS-CoV-2 reactive memory T cells in convalescent patients were more prevalent to antigens that are not subject to mutational variation such as the N protein and Le Bert *et al* demonstrated that N protein specific T cell responses induced by SARS-Co-V were more likely to react with homologous SARS-CoV-2 peptides than those from NSP7 and NSP13 ^18,44,45^. This supports the rationale for designing vaccines to induce T cell and antibody mediated immunity to both S and N proteins, such as the strategy described in this study.

Herein, we have also achieved simultaneous production of strong neutralising antibody titres to both SARS-CoV-2 S and N proteins. The RBD monomer elicited the strongest neutralising antibody responses with ID_50_ titres of >5000. In this study, these titres were higher than obtained with the NIBSC 20/136 reference control and those elicited by a DNA vaccine encoding whole S protein, which was also reflected in the ACE2 binding inhibition assay, suggesting that vaccination with RBD may elicit higher neutralising antibody titres than vaccination with the complete S antigen. Despite the fact that this may be linked to the protein expression levels from the vaccines in this study, these findings are concordant with other studies in mice in which the whole S protein or an RBD trimer have been delivered as recombinant protein, RNA or adenovirus vectors in prime boost regimes ^19,46–51^. Our *in vivo* efficacy data demonstrate that the antibody titres and T cell responses induced were sufficient to protect mice from lethal viral challenge and promote survival. It has been demonstrated that the majority of the neutralising antibodies to S protein target the RBD and as well as high levels of protein expression, support the inclusion of just the RBD sequence in this bivalent vaccine ^52^.

The inclusion of the N protein sequence in this vaccine was not only intended for the stimulation of T cell responses; antibody responses to the N protein were also detected. Although, these antibodies do not prevent virus acquisition, N specific antibodies can often clear viral infections ^53–55^. Reports using lymphocytic choriomeningitis virus (LCMV) suggest an alternative role of N protein specific antibodies that involves antibody-dependent intracellular neutralisation and promotion of N specific cytotoxic T lymphocytes (CTLs) that efficiently clear virus infected cells ^56^. Caddy *et al*. suggest that N-specific antibodies bind N protein released during viral lysis or expressed on the surface of infected cells and that immune complexes can be taken up by APCs, in which TRIM21 targets N protein for cytosolic degradation and the generation of cytotoxic T cells. This mediates viral clearance prior to the establishment of neutralising antibodies. The vaccine construct containing N protein fused to the modified Fc sequence induced significantly better T-cell responses to N protein and elicited higher responses to RBD than a similar construct expressing the same RBD construct, but with unmodified Fc linked to N. This suggests that the modified N-Fc fusion is enhancing the T-cell response to other antigens and perhaps the N specific antibodies could play a role in viral clearance and protection. A recent report by Matchett et al. has demonstrated the protective ability of an adenovirus type-5 (Ad5)-vectored vaccine encoding N protein from SARS-CoV-2 in murine and hamster infection models and shows that protection is associated with the rapid stimulation of N protein specific T cell responses ^57^, thereby providing further evidence for the important role of N protein specific T cell responses. The role of anti-N protein immunity induced by the vaccine described in this report would need to be determined in future studies in the absence of the RBD sequence. Reports by Rice *et al* and Seiling *et al* have also demonstrated the benefit of combining both S and N proteins as vaccine targets using a Ad5 vectored delivery platform ^19,58^.

The emergence of new SARS-CoV-2 variants has highlighted the importance of generating a vaccine mediated response that is effective against the variants as well as the original strain. Mutations in the S protein have been shown to reduce antibody recognition and neutralisation and increase the transmissibility, infection rate and disease severity ^15,16,59,60^. Approved vaccines only encode the original Lineage A S protein sequence and have shown varying levels of cross-reactivity against variant S proteins, with most reporting a drop in recognition and neutralisation efficacy, particularly against the Beta variant ^52,61,62^. In other reports, higher cross-reactivity is retained against the Alpha variant, which has fewer significant mutations within the RBD sequence ^60,63^. Next generation vaccines now cover sequences of virus variants including Omicron. In our studies we have demonstrated a decrease in antibody titres, ACE2 binding inhibition and pseudotype neutralisation against the Beta and Gamma variants in sera from mice immunised with the Lineage A vaccine with activity against the Alpha and Delta variants being largely maintained. Responses remain superior to those seen in the sera from mice immunised with whole S encoding DNA and in the NIBSC 20/136 reference control serum.

We have also demonstrated the efficacy of a vaccine construct encoding the Beta variant RBD and show high titre antibody responses to the Beta variant S protein that cross reacts with the Lineage A S protein. Reports suggest that sera from patients with the Beta variant infection have high titre cross neutralising antibodies that also recognise the Lineage A SARS-CoV-2 virus ^64^. In this report we demonstrate that although sera from mice immunised with the Beta variant vaccine show a reduction in ACE2 binding inhibition of the Lineage A and Alpha variants, responses remain above 50% inhibition against all variants tested and show higher neutralisation titres than to the NIBSC 20/136 control that are also comparable with those induced by current vaccines targeting S protein only ^52,65^. At the time of this study the Beta variant was the greatest concern and these data imply that there is evidence of cross-reactivity therefore use of a vaccine encoding the Beta variant sequences may provide additional protection against multiple SARS-CoV-2 variants. Such variant vaccines have potential to be used to boost waning immunity.

DNA provides a cost effective and rapid to manufacture product that can be easily adapted to encode sequences or mutations present in new variants. DNA vectors also lend themselves well to use in multiple homologous prime boost immunisation regimes, which is often not possible with other viral vector-based vaccination strategies due to limitations of anti-vector immunity but may be vital to combat rapidly emerging virus variants. Data from our cancer vaccine platform demonstrates the stability of the DNA vector used in this study and its safety in human subjects ^23^. DNA does have limitations, in particular the apparent low immunogenicity in humans, possibly a reflection of limitations in cellular uptake and nuclear delivery for efficient protein expression. Electroporation has been shown to be beneficial to enhance this, although this delivery method is not as practical for prophylactic vaccine rollout due to the requirement for specialised equipment. Consistent with data from others, unpublished data with our SCIB1 vaccine using the same vector backbone has suggested negligible uptake and persistence of the DNA in tissues or organs distal to the immunisation site indicating a low risk of genomic integration. In comparison to the ZyCov-D and INO-4800 vaccines encoding the S protein, the data described herein demonstrate stronger antibody (IgG end point and neutralisation titres) and T cell responses from a 10-100 times lower DNA dose in mouse models ^49,66^. However, it is difficult to directly compare data as this report details the delivery of DNA using the gene gun and not direct needle injection or electroporation. However, our unpublished data in rats with DNA delivery via a needle-free device compares favourably with similar delivery and dose of the ZyCoV-D data in rabbits showing higher antibody titres and T cell responses. The needle free delivery devices are being used in clinical studies to deliver DNA vaccines and this vaccine is also currently in clinical trials using one such needle free delivery device. The bivalent DNA SARS-CoV-2 vaccine in this study stimulates strong humoral and cellular immunity that can be easily adapted to emerging virus variants. Furthermore, the responses elicited by both Lineage A and Beta variant vaccines show cross-reactivity with multiple viral variants and support the potential clinical application of these vaccines.

## Methods

### DNA plasmids

The backbone of the vaccine plasmids was derived from the FDA regulatory compliant vector backbone of pVAX1 (Invitrogen) for use in humans. All nucleotide sections for insertion were codon optimised for expression in humans and contain a human IgH leader (MDWIWRILFLVGAATGAHS). Codon-optimised nucleotide sections encoding the leader, amino acids (aa) of the S glycoprotein RBD domain 319–541 or 330–525 (GenBank accession number YP_009724390, isolate Wuhan-Hu-1) alone, fused in frame with either the Hinge-CH2-CH3 domain of the human IgG1 Fc constant region (GenBank accession number P01857) or the variant Hinge-CH2-CH3 iV1 (where 23 amino acids (aa) have been replaced with murine IgG3 residues) or attached to a fibritin trimer fold on (GYIPEAPRDGQAYVRKDGEWVLLSTFL) or disulphide bridge motif (CCGGGSG) via a glycine serine linker were synthesised with *BamH*I and *Xho*I sites inserted at the 5’ and 3’ends respectively. In the first round of cloning, these sections were inserted into the *BamH*I/*Xho*I sites of the pVaxDCIB68 (SCIB1) plasmid as described previously ^20,67^, in direct replacement of the SCIB1 light kappa chain in the first expression cassette to generate intermediate plasmids.

In a second round of cloning codon-optimised nucleotide sections encoding the leader, fulllength nucleoprotein aa 2–419 (Accession number YP_009724397) alone or fused in frame with the Hinge-CH2-CH3 domain of the human IgG1 constant domain or the variant Hinge-CH2-CH3iV1 were synthesised and flanked with *Hind*III/*Pst*I. The heavy chain was excised using *Hind*III/*Pst*I from the intermediate vectors generated from the first round and replaced with the N protein sections in the second expression cassette alongside the appropriate S section (Figure 1A). The sequences of both chains within each expression cassette of the pVaxDC vectors were confirmed by the dideoxy chain termination method ^68^.

The plasmid pCMV3-2019-nCoV-Spike (S1+S2)-long encodes full-length spike from SARS-CoV-2 amino acid 1–1273 (GenBank accession number YP_009724390 /QHD43416.1) and was obtained from Sino Biological (catalogue number VG40589-UT). This contained codon optimised cDNA for expression of the protein in mammalian cells inserted into the *Kpn*I/*Xba*I sites of the mammalian expression vector pCMV3-untagged under control of the high-level expression mammalian human enhanced cytomegalovirus (CMV) immediate early promoter.

### Transient Expi293F™ transfection

Secretion levels from plasmid DNA constructs were evaluated following transient transfections of Expi293F™ cells using the ExpiFectamine™ 293 Transfection kit (Gibco, LifeTechnologies). Briefly, Expi293F™ cells in suspension (100 mL, 2×10^6^/mL) were transfected with 100μg DNA and conditioned medium harvested at day 6 post-transfection. Conditioned supernatant was filtered through 0.22 μm bottle top filters (Merck Millipore) and sodium azide added to a final concentration of 0.2% (w/v). Cell pellets were stored at −80°C. Cell lysates (intracellular RBD and N protein) were generated by processing the cell pellets in a suitable volume of RIPA buffer (Sigma Aldrich cat# R0278) according to the manufacturer’s recommendations.

### Sandwich ELISA for detection of secreted and intracellular RBD, S and N protein

Commercial kits/antibody pairs were used in both cases. N protein was detected using the SARS-CoV-2 NP ELISA kit from Bioss (cat# BSKV0001) according to the supplier’s instructions. Quantitation relied on the standard curve generated using the N standard supplied with the kit. RBD was detected using a sandwich ELISA consisting of a capture antibody (SARS-CoV-2 S neutralising mouse monoclonal antibody (mAb) Sino Biological, cat# 40591-MM43) combined with an HRPO-labelled detection antibody from the SARS-CoV-2 spike RBD Antibody Pair (Epigentek cat# A73682). Capture antibody was coated at 200 ng/well and detection antibody was used at a dilution of 1:1000. S protein was detected using a pre-coated sandwich ELISA from ACRO Biosystems with biotinylated anti-spike protein detection antibody and streptavidin-HRP (ACRO Biosystems cat# RAS-A020).

### Peptides and proteins

Peptides were selected based on immune epitope database (IEDB) (http://www.iedb.org/) binding predictions ^69^ and from reports in the literature. Peptides (Table S1) were synthesised at >90% purity (GenScript), stored lyophilised at −80°C and then reconstituted in phosphate buffered saline (PBS) on day of use. SARS-CoV-2 RBD and N protein peptide pools were purchased from JPT Peptide Technologies and GenScript, respectively. SARS-CoV-2 N and S1 proteins were obtained from Genscript and Sino Biological.

### Animals and immunisation protocol

Animal experiments were carried out with ethical approval under a UK Home Office-approved Project License. HLA-A2 (HLA-A2.1+/+ HLADP4+/+ hCD4+/+ (HHDII/DP4; EM:02221, European Mouse Mutant Archive), HLA-A2/DR1 (HHDII/DR1; Pasteur Institute), HLA-A2 (HHDII; Pasteur Institute)) transgenic mice or BALB/c and C57Bl/6 (Charles River) aged 7–12 weeks were used. CD rats (Charles River) were used at 225-250g weight were used.

Mice or rats were immunised with 1 μg plasmid DNA, coated onto gold particles, intradermally using a Helios™ gene gun (BioRad, Hemel Hempstead, UK) on days 1, 8 and 15 and responses analysed on day 21, unless stated otherwise.

### ELISpot assays

Murine and rat ELISpot kits (Mabtech) were used with 5×10^5^ splenocytes/well in RPMI with L-glutamine, 10% (v/v) foetal bovine serum (FCS) supplemented with penicillin, streptomycin, HEPES and 10^-5^M 2-β mercaptoethanol (Invitrogen). Synthetic peptide (5 μg/mL) or peptide pool (1 μg/mL) were added to triplicate wells. CD8, or CD4 antibodies (Supplementary table 2) were added for blocking studies (20 μg/mL). Plates were incubated at 37°C for 40 hours in an atmosphere of 5% (v/v) CO_2_. Spots were counted using an automated plate reader (Cellular Technologies Ltd).

### Cytokine analysis using Mesoscale Discovery (MSD) platform

Supernatant from *ex vivo* IFNγ ELISpot assays were collected and analysed on the MSD platform using a murine U-PLEX custom biomarker kit according to manufacturer’s instructions. In brief, analyte in culture supernatant is captured onto the plate by specific antibodies and detection antibodies conjugated with electrochemiluminescent labels bind to the analytes to complete the sandwich immunoassay. A voltage applied to the plate electrodes causes the captured labels to emit light which provides a quantitative measure of each analyte in the sample.

### Intracellular cytokine analysis

Splenocytes (5×10^6^/mL) were incubated with 1 μg/mL synthetic RBD and N peptide pools or 0.5 μg/mL whole recombinant S and N proteins for 7 days. Cells were washed, plated at 5×10^6^/mL and stimulated for 8–12 hr with medium alone, 0.25 μg/mL CD3 mAb (clone 145-2C11, ThermoFisher) as a positive control, 5 μg/mL synthetic peptide or 1 μg/mL peptide pool. Cells were incubated at 37°C for 8 hr with Brefeldin A (BD Biosciences) and Monensin (ThermoFisher) being added for the last 7 hr of incubation. After 8 hr incubation, cells were held at 4°C until staining. Cells were then washed and stained with APC efluor™ 780 conjugated anti-mouse CD4 and VioGreen™ conjugated anti-mouse CD8 mAbs. Cells were subsequently washed, fixed and permeabilised using intracellular fixation/permeabilisation buffers (ThermoFisher). Intracellular antibody staining was performed in permeabilisation buffer for 30 min before washing and fixing. Stained samples were analysed using a Milteny Biotech MACSQuant™ 10 or 16 flow cytometer. Full details of the antibodies used are provided in Table S2.

### ELISA for N, S1 and RBD specific antibodies

Blood collected from mice was allowed to clot for at least 30 min at room temperature, centrifuged at 1000g for 10 min and serum collected. Serum from naïve mice was used as a control. Sera were diluted in 2% (w/v) bovine serum albumin in PBS (BSA-PBS) and incubated for 1 hr at room temperature (RT) in coated (S1 antigen, RBD antigen or N-protein, at 200 ng/well, Supplementary Table 1) and blocked ELISA plates. Antibody binding to S1 and RBD antigen was detected using a three-step approach consisting of anti-mouse or antirat Fc biotin (Sigma Aldrich B7401) followed by streptavidin-HRPO (Life Technologies cat# SA1007) and finally TMB (3,3’, 5, 5’-tetramethylbenzidine) Core+ reagent (Bio-Rad cat # BUF062C) for development. Antibody binding to N protein was detected using conventional two-step ELISA with an HRPO-conjugated anti-mouse or anti-rat Fc antibody (Sigma Aldrich cat # A9309). All assays included a standard curve of mouse neutralising S1 antibody and mouse anti-N antibody (Sino Biological). Standard curves were used to extrapolate the amount (μg/mL) of specific immunoglobulin (Ig) in the sera samples.

### Pseudotype neutralisation assay

SARS-CoV-2 S protein plasmids were generated and cloned, and pseudoparticles generated as previously described ^70,71^. Briefly, 1.5 × 10^6^ HEK293T cells were seeded overnight in a 10 cm diameter Primaria-coated dish (Corning). Transfections were performed with 2 μg each of the murine leukemia virus (MLV) Gag-Pol packaging vector (phCMV-5349), luciferase encoding reporter plasmid (pTG126) and plasmid encoding Wuhan strain SARS-CoV-2 S using 24 μL polyethylenimine (Polysciences) in Optimem (Gibco). Reaction mixtures were replaced with Complete Dulbecco’s Modified Eagle Medium growth medium after 6 hr. A no-envelope control (empty pseudotype) was used as a negative control in all experiments. Supernatants containing SARS-CoV-2 pseudotype viruses were harvested at 72 hr posttransfection and filtered through 0.45 μm membranes. Pseudotypes generated in the absence of S were used as a negative control. For infectivity and neutralisation testing of SARS-CoV-2 pseudoparticles, 1 × 10^5^ VeroE6 cells were seeded on white 96-well tissue culture plates (Corning) and incubated overnight at 37°C. The following day, SARS-CoV-2 pseudotypes were mixed with appropriate amounts of heat inactivated serially diluted sera and then incubated for 1hr at RT before adding to cells. Naïve mouse serum was tested at 1/50 dilution as a control. After 72 hrs at 37°C, cells were lysed with cell lysis buffer (Promega cat # E1500) and placed on a rocker for 15 min. Luciferase activity was then measured in relative light units (RLUs) using a FLUOstar Omega plate reader (BMG Labtech) and MARSdata analysis software. Each sample was tested in triplicate. Neutralising activities were reported as reciprocal serum dilution levels corresponding to 50% inhibitory dilution (ID50) values and were calculated by nonlinear regression (GraphPad Prism version 9.1.2), using lower and upper bounds (0% and 100% inhibition) as constraints to assist curve fitting.

### Live virus neutralisation assay

SARS-CoV-2 infectious virus (CVR-GLA-1) was obtained from the Centre for AIDS Reagents, NIBSC, UK. Live virus neutralisation assays were performed using method previously described ^72^, except that 790 TCID_50_/mL of the SARS-CoV-2 virus was added to each serum dilution. Additionally, for some experiments, the sera were diluted down to 1:81,920.

### ACE2 binding inhibition assay

A V-PLEX COVID-19 ACE2 neutralisation kit from Meso Scale Diagnostics LLC was used to investigate the ability of vaccine-elicited antibodies to block the binding of ACE2 to RBD or whole S proteins. V-plex SARS-CoV-2 Panel 11 and 13 multispot plates containing S1 RBD and whole S proteins respectively from Lineage A (originally identified in Wuhan) and variant (B1.1.7/Alpha, B1.351/Beta, P.1/Gamma, B.1.617.2/Delta) SARS-CoV-2 strains were blocked, followed by incubation with sera at 1:100 dilution and Sulfo-tagged human ACE2 protein, according to the manufacturer’s instructions. Results are expressed as percentage inhibition of ACE2 binding via comparison of sera-incubated samples to diluent-containing wells (absence of inhibition).

### In vivo virus challenge model

SARS-CoV-2 viral challenge experiment ^73^ was performed at Texas Biomedical Research Institute (Texas Biomed; San Antonio, Texas, USA). Texas Biomed Biosafety and Institutional Animal Care and Use Committees approvals (IACUC) were obtained prior to conducting experiments. The infected animals were housed under Animal Biosafety Level 3 (ABSL3) facilities at the Southwest National Primate Research Center, where they were treated according to the standards recommended by AAALAC International and the NIH Guide for the Care and Use of Laboratory Animals. K18 hACE2 transgenic mice (Jackson Laboratories) were immunised with DNA via gene gun on days 0 and 28 followed by infection intranasally (i.n.) with 1×10^4^ plaque-forming units (PFU) of SARS-CoV-2 (USA-WA1/2020 strain) in a final volume of 50 μL following isoflurane sedation on day 56. Unimmunised and unchallenged (PBS control) mice were used as controls. Body weights and survival were monitored over time. Nasal turbinates and lungs were assessed on day 58 for viral load via plaque assay ^73^.

### Statistical analysis

Statistical analysis was performed using GraphPad Prism software version 9. Comparative analysis of the ELISpot results was performed by applying paired or unpaired ANOVA as appropriate with Sidak’s multiple comparisons test and P values calculated accordingly. P<0.05 values were considered statistically significant.

## Acknowledgments

Work was funded by Scancell Ltd and grant from UKRI. We would like to disclose that a number of authors are employed by Scancell Limited. L.G.D and S.E.A are directors and shareholders of Scancell Ltd, and A.G.P is CEO of Multimmune GmbH. M.A, J.D, V.D and V.K are employees of Texas Biomedical research Institute. The manuscript has been reviewed by Dr T Parsons.

## Author Contributions

Conceptualization – L.G.D, J.K.B, J.M.D, S.E.A, Formal Analysis – V.A.B, M.V, Funding acquisition – L.G.D, S.E.A, J.E.D, A.G.P, J.M.D, J.K.B, Investigation – V.A.B, M.V, R.L.M, P.S, K. W.C, R.A.U, T.T, C.M.C, K-C.C, A.S, E.D, I.D, S.S, M.A, J.D, V.D, V.K, Project administration – L.G.D, S.P, V.A.B, Resources – R.L.M, A.G.P, J.E.D, Supervision – L.G.D, V.A.B, M.V, Validation – S.P, S.E.A, Visualization – V.A.B, S.P, Writing (original draft) – V.A.B, L. G.D, Writing (review & editing) – V.A.B, L.G.D, M.V, R.A.U, S.E.A, S.J.P, A.G.P, C.M.C, J.M.D, J.K.B, V.K.

