## Supplementary figures and tables for "SARS-CoV-2 spike RBD and nucleocapsid encoding DNA vaccine elicits T cell and neutralising antibody responses that cross react with variants"

### Supplementary Table 1. Peptides and Proteins

| Antigen | Peptide/protein coordinates | Supplier (cat. no.) | Predicted binding (IEDB) | | | | | | | | ref |
|---|---|---|---|---|---|---|---|---|---|---|---|
| | | | H-2Kd | H-2Dd | H-2Kb | H-2Db | I-Ab | I-Ad | I-Ed | HLA-A2 | |
| Spike | Overlapping pool | JPT (PM-WCPV-S-RBD-1) | | | | | | | | | |
| | 417-425 | Genscript, (custom) | 2.4 | 0.89 | 0.72 | 10 | | | | 0.05 | [36] |
| | 505-524 | Genscript, (custom) | 0.56 | 0.04 | 0.01 | 0.17 | 4.65 | 18.0 | 14.6 | 0.45 | [52] |
| | S1 protein (Lineage A) | Genscript (Z03501) | | | | | | | | | |
| | S1 protein (B.1.1.7) | Sino Biological (40591-V08H12) | | | | | | | | | |
| | S1 protein (B.1.351) | Sino Biological (40591-V08H10) | | | | | | | | | |
| | RBD protein (B.1.351) | Sino Biological (40592-V08H90) | | | | | | | | | |
| | RBD protein (B.1.617.2) | Sino Biological (40592-V08H85) | | | | | | | | | |
| | S1 protein (B.1.1.529) | Genscript (Z03729) | | | | | | | | | |
| Nucleoprotein | Overlapping pool | Genscript (RP30013) | | | | | | | | | |
| | 80-100 | Genscript, (custom) | 0.14 | 4.2 | 2.1 | 12 | 41.0 | 30.5 | 0.1 | 13 | |
| | 101-121 | Genscript, (custom) | 0.38 | 0.3 | 3.3 | 2.6 | 1.24 | 67.0 | 8.6 | 5.3 | |
| | 138-146 | Genscript, (custom) | 1.4 | 5.6 | 11 | 1.9 | | | | 1.7 | [35] |
| | 212-231 | Genscript, (custom) | 2.4 | 1.7 | 0.61 | 0.19 | 50.0 | 45.5 | 42.0 | 0.02 | |
| | 299-318 | Genscript, (custom) | 0.35 | 0.03 | 0.21 | 0.18 | 0.33 | 8.65 | 35.5 | 0.91 | |
| | 301-321 | Genscript, (custom) | 0.35 | 0.03 | 0.21 | 0.18 | 0.33 | 12.3 | 7.1 | 0.91 | |
| | protein | Genscript (Z03488) | | | | | | | | | |

Supplementary Table 2. Antibodies

| **Antibody target** | **Target species** | **Format** | **Clone** | **Supplier** |
|---|---|---|---|---|
| CD8a | mouse | VioGreen™ | REA601 | Miltenyi |
| CD4 | Mouse | APC-eFluor™780 | RM4-5 | Thermofisher |
| CD8 | Mouse | Functional grade | 2.43 | BioXcell |
| CD4 | Mouse | Functional grade | GK1.5 | BioXcell |
| IFNγ | Mouse | PE-Vio™770 | XMG1.2 | Thermofisher |
| TNFα | Mouse | FITC or APC | MP6-XT22 | Thermofisher |
| IL-4 | Mouse | APC | 11B11 | Thermofisher |
| IL-10 | Mouse | Pacific Blue™ | JES5-16E3 | Biolegend |
| Spike S1 | SARS-CoV-2 | purified | MM43 | Sino Biological |
| Nucleocapsid | SARS-CoV-2 | purified | MM05 | Sino Biological |

Supplementary Table 3. Mutations with RBD and N protein sequences of variants of concern

| Pango lineage | **B.1.1.7** | **B.1.351** | **P.1** | **B.1.617.2** | **B.1.1.529** |
|---|---|---|---|---|---|
| WHO label | Alpha | Beta | Gamma | Delta | Omicron |
| VoC or VuI | VOC-20DEC-18 | VOC-20DEC-18 | VOC-21JAN-11 | VOC-21MAY-11 | VOC-21NOV-26 |
| First identified | Kent, UK | South Africa | Brazil | India | Multiple countries |
| Mutations reported in Spike RBD (319-541) | N501Y | K417N<br>E484K<br>N501Y | K417N<br>E484K<br>N501Y | L425R<br>T478K | G339D<br>S371L<br>S373P<br>S375F<br>K417N<br>N440K<br>G446S<br>S477N<br>T478K<br>E484A<br>Q493R<br>G496S<br>Q498R<br>N501Y<br>Y505H |
| Mutations reported in N protein | D3L<br>S235F | T205I | P80R | D63G<br>R203M<br>D377Y | G204R<br>R203K |

Supplementary Figure 1

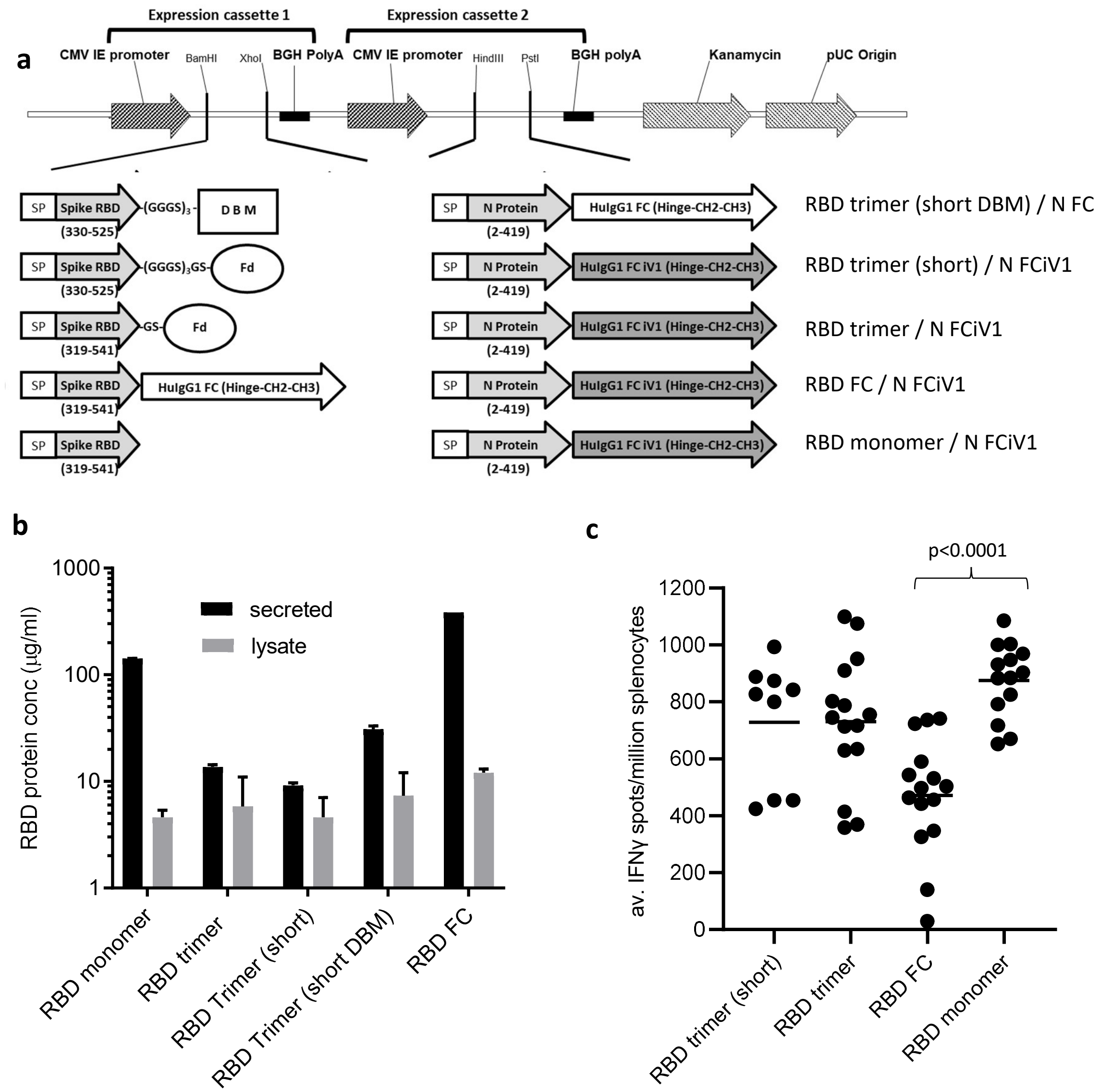


Figure S1. A, Schematic representation of the DNA SARS-CoV-2 plasmids and expression of RBD proteins. A, DNA constructs expressing RBD variants. Spike RBD and NP variant chains are within expression cassettes 1 and 2 of the pVaxDC vector. Numbers indicate amino acids. SP: signal peptide/ Human IgH Leader, RBD: receptor-binding domain, Fd: Fibritin fold on trimer motif from T4 bacteriophage (GYIPEAPRDGQAYVRKDGEWVLLSTFL). DBM: Disulphide bridge motif. Both fibritin fold on and disulphide bridge motif are attached to the S RBD via glycine/serine linkers. In some constructs N protein is fused inframe with HuigG1 FC or with the improved modified version of the constant domain, designated iV1.
B, RBD protein concentration in supernatant or lysate of Expi293F™ cells transfected with RBD trimer variants measured by RBD protein ELISA. C, RBD peptide pool specific T cell responses in mice immunised with DNA constructs expressing RBD variants alongside N linked to modified Fc via gene gun on days 1, 8 and 15. Responses measured by IFNγ ELISpot assay and normalised against unstimulated control. Data is collated from at least 3 independent studies in which n=3.

Supplementary Figure 2

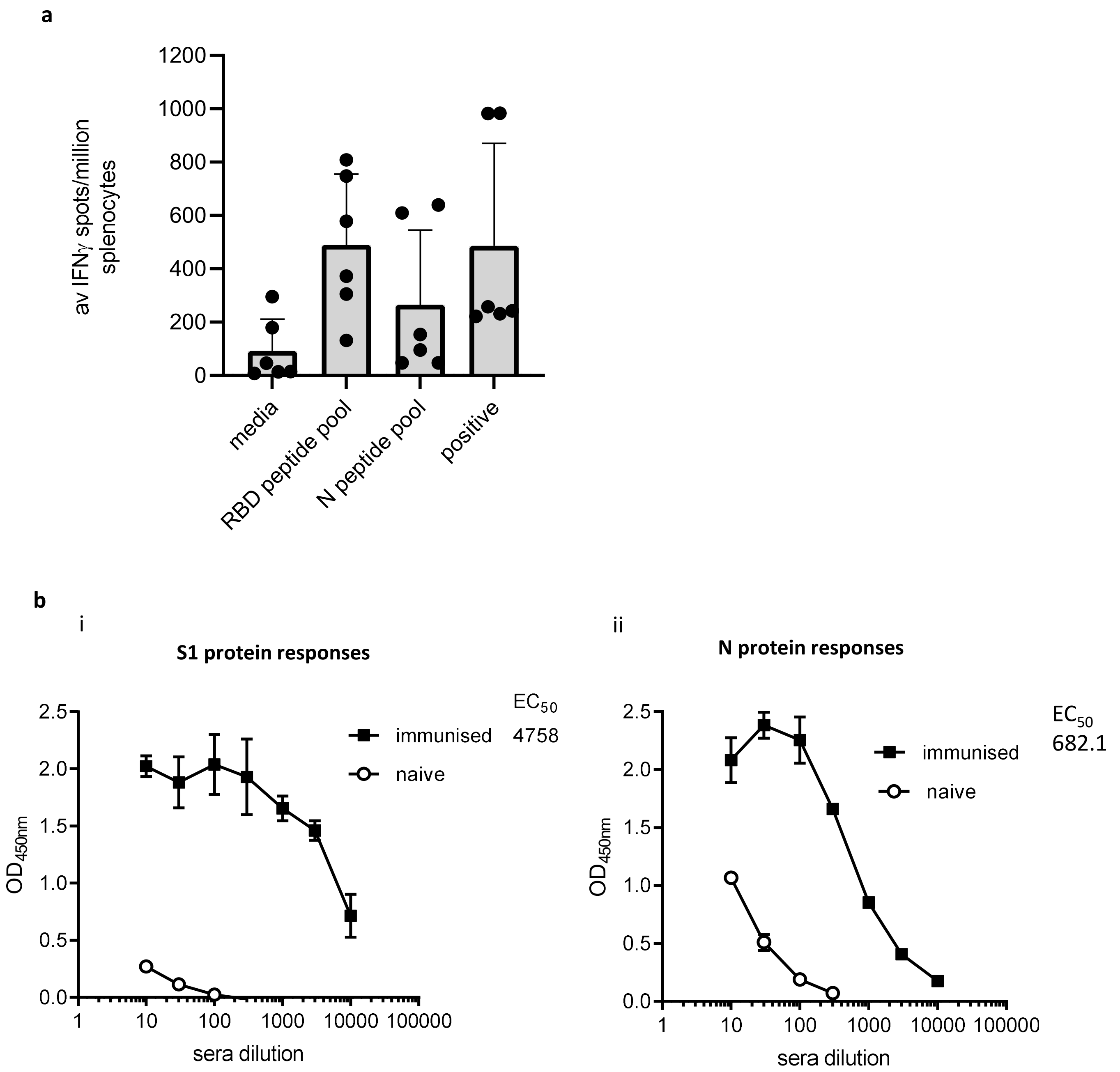


Figure S2. Vaccine induced responses in rats. CD rats immunised with DNA construct expressing RBD monomer and N linked to modified Fc (iV1) via gene gun on days 1, 8 and 15 or days 1 and 29. A, T cell responses monitored by IFNγ ELISpot assay to RBD and N peptide pools. Symbols represent mean response for individual rats, line/bar represents mean value between rats. B, Antibody responses to S1 (i) and N (ii) proteins measured by ELISA in sera from immunized rats compared to naïve sera.
Data is collated from at least 2 independent studies in which n=3.

### Supplementary Figure 3

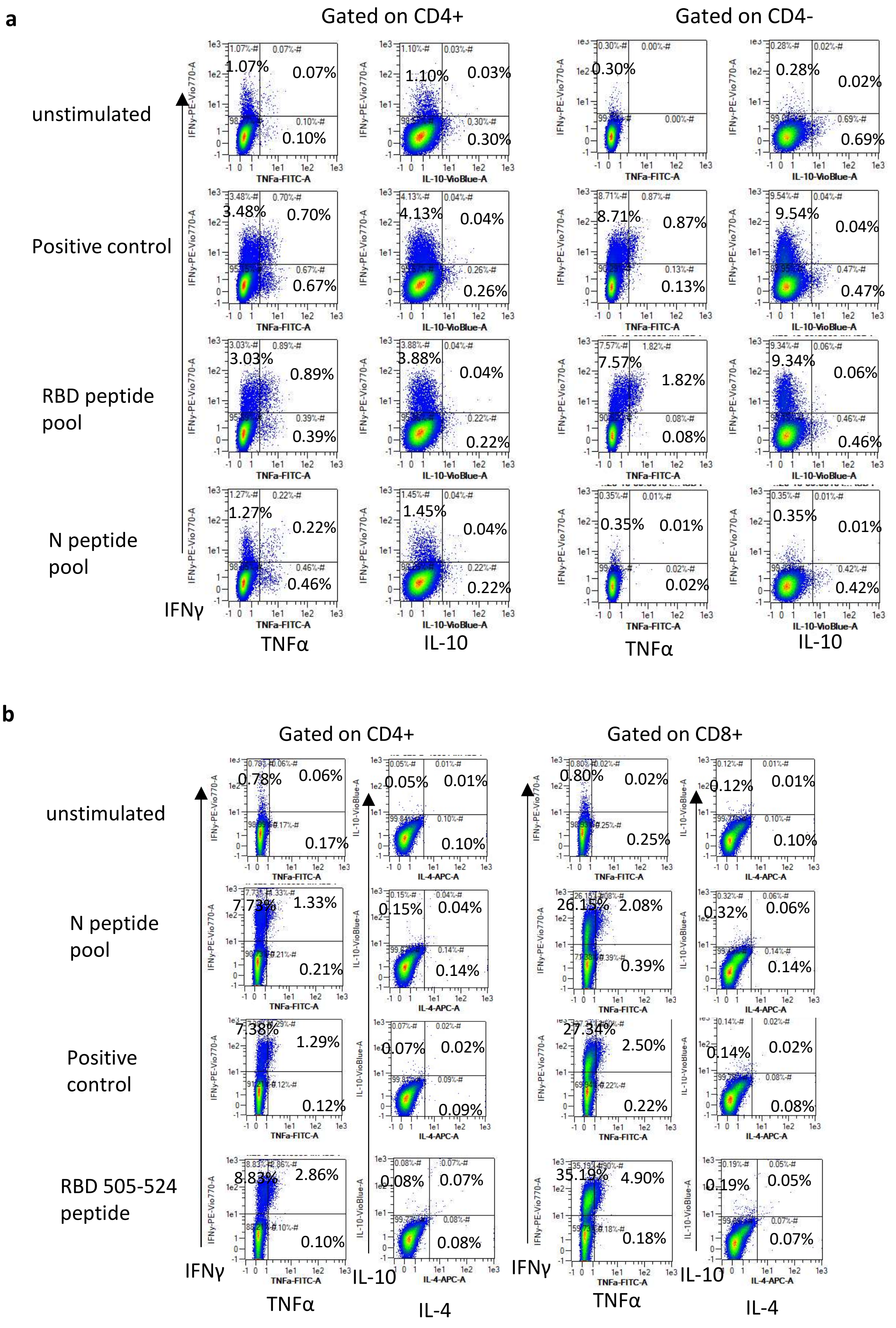


Figure S3. Multi-cytokine T cell responses with Th1 polarisation. Example characterization and intracellular cytokine staining for IFNγ, TNFα, IL-10 and IL-4 in response to peptide restimulation or stimulation with anti-CD3 antibody positive control on splenocyte cultures expanded in vitro with S1 and N proteins or peptide pools from BALB/c (A) or C57Bl/6 (B) mice immunised with DNA construct expressing RBD alongside N linked to modified Fc (iV1). Percentage of CD4+ or CD8+ cells are shown for responses.

Supplementary Figure 4

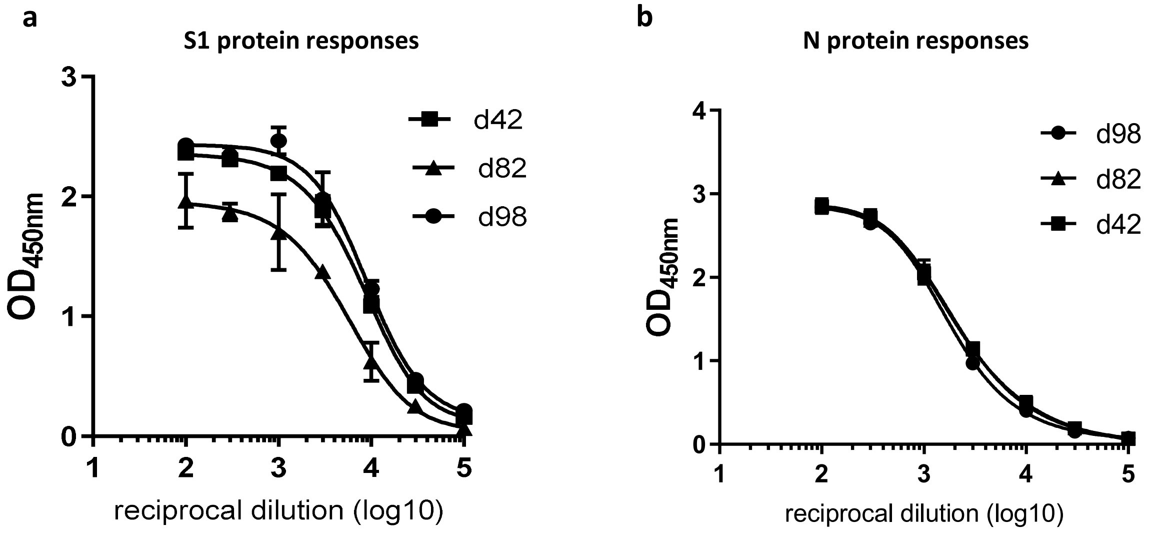


Figure S4. BALB/c mice immunised with DNA construct expressing RBD monomer and N linked to modified Fc via gene gun on days 1, 29 and 85 and sera analysed at days 42, 82 and 98. Antibody responses to S1 (A) and N (B) proteins measured by ELISA.
Data is representative of at least 2 independent studies in which n=3.

Supplementary Figure 5

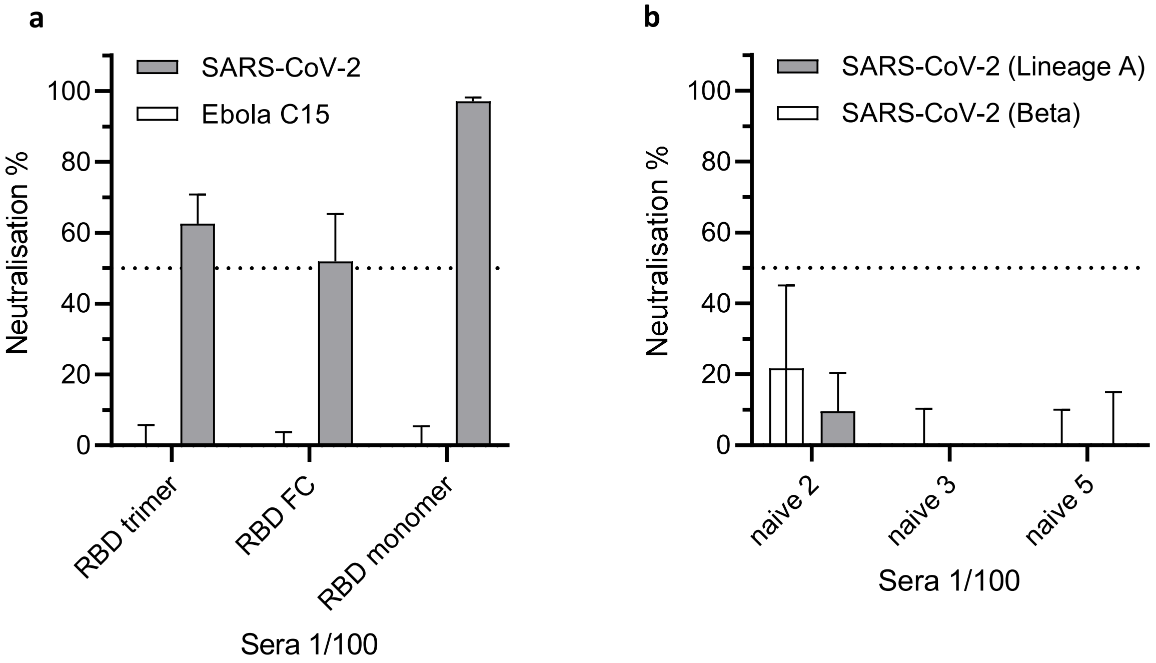


Figure S5. A, Pseudotype neutralisation assay of sera from mice with DNA expressing RBD trimer, monomer or linked to FC against SARS-CoV-2 or irrelevant Ebola pseudotypes. B, Pseudotype neutralisation of serum from naïve mice against SARS-CoV-2 lineage A or Beta strains. Assays performed with serum at 1/100 dilution and is representative of at least two independent experiments.

Supplementary Figure 6

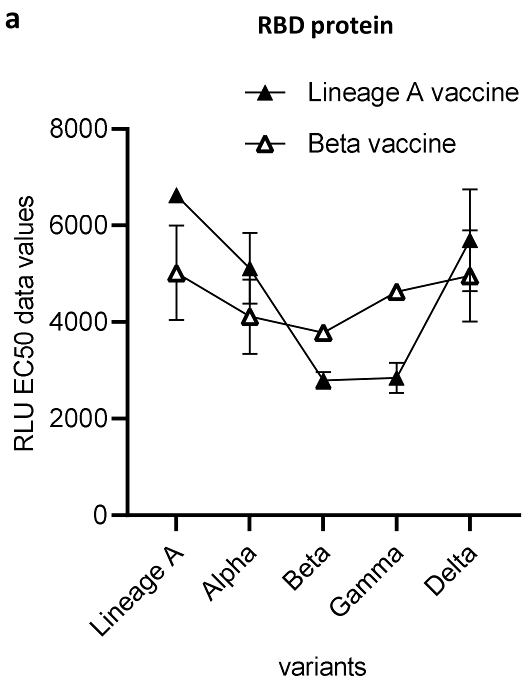


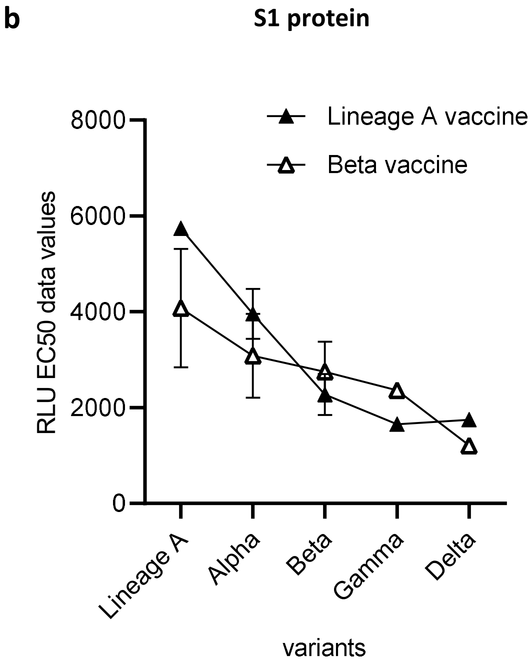


Figure S6. Antibody responses to variant RBD (A) or whole spike (B) proteins measured by MesoScale Discovery assay in sera from mice immunised with Lineage A or Beta variant vaccines. Data plotted for average sera dilution EC50 values and shown with 95% Confidence Interval as a measure of variation.

Supplementary Figure 7

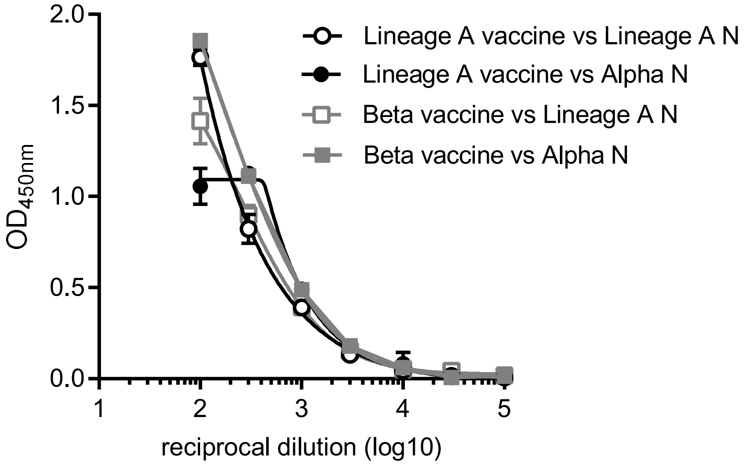


Figure S7. Antibody responses to variant N proteins measured by ELISA in sera from mice immunised with Lineage A or Beta variant vaccines

Supplementary Figure 8

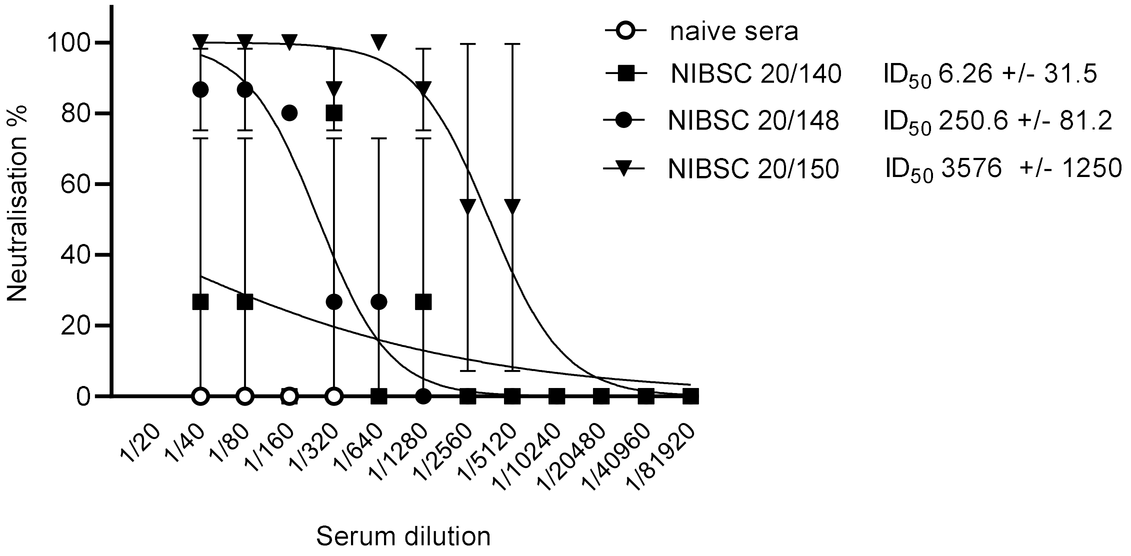


Figure S8. Live virus neutralisation assay using NIBSC reference standards. $IC_{50}$ values were calculated and shown with 95% Confidence Interval as a measure of variation.

Supplementary Figure 9

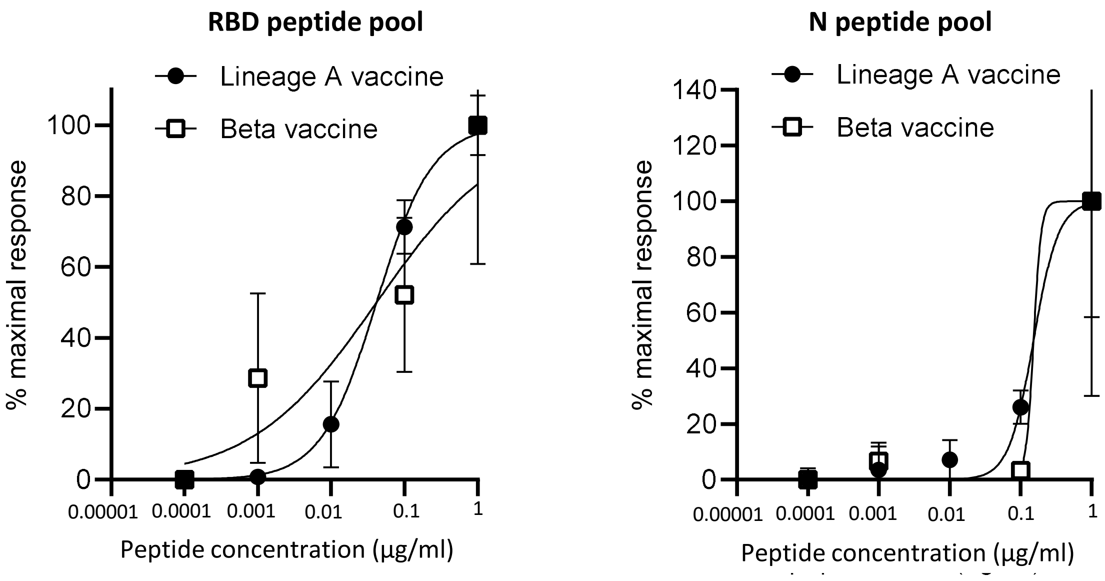


Figure S9. RBD and N peptide pool response titration. Mice immunized with DNA construct expressing Lineage A or Beta RBD monomer and N linked to modified Fc (iV1) via gene gun on days 1, 8 and 15. T cell responses monitored by IFNγ ELISpot assay to titrating amounts of (A) RBD and (B) N peptide pools. Data normalized against unstimulated (media) control and each point is an average from at least 2 individuals. Data is representative of at least 2 independent studies.

.
